# Dietary restriction improves fitness of ageing parents but reduces fitness of their offspring

**DOI:** 10.1101/632026

**Authors:** Brian S. Mautz, Martin I. Lind, Alexei A. Maklakov

**Affiliations:** Animal Ecology, Department of Ecology and Genetics, Uppsala University, Norbyvägen 18D, Uppsala 752 36, Sweden; Division of Epidemiology, Department of Medicine, Vanderbilt University Medical Center, Nashville, TN 37203, USA; School of Biological Sciences, University of East Anglia, Norwich Research Park, Norwich NR4 7TJ, Norfolk, UK

## Abstract

Dietary restriction (DR) is a well-established intervention to extend lifespan across taxa. Recent studies suggest that DR-driven lifespan extension can be cost-free, calling into question a central tenant of the evolutionary theory of ageing. Nevertheless, boosting parental longevity can reduce offspring fitness. Such intergenerational trade-offs are often ignored but can account for the ‘missing costs’ of longevity. Here, we use the nematode *Caenorhabditis remanei* to test for effects of DR by fasting on fitness of females and their offspring. Females deprived of food for six days indeed had increased fecundity, survival and stress resistance after re-exposure to food compared to their counterparts with constant food access. However, offspring of DR mothers had reduced early and lifetime fecundity, slower growth rate, and smaller body size at sexual maturity. These findings support the direct trade-off between investment in soma and gametes challenging the hypothesis that increased somatic maintenance and impaired reproduction can be decoupled.

## Introduction

Dietary restriction (DR), defined as a reduction in nutrient intake without malnutrition, is known to increase lifespan across the tree of life, from unicellular organisms to worms to rodents to primates^1–4^. Moreover, DR is often associated with improved stress resistance^5^, reduced rate of ageing^6^ and even improved reproduction in late life^7^. Despite the ubiquity of DR-driven lifespan extension, and considerable multidisciplinary interest in this topic, the evolutionary underpinnings of this effect remain poorly understood. The leading evolutionary hypothesis maintains that DR is an evolved resource allocation strategy aimed at increasing survival during the time of temporary famine^8^. This “waiting-for-good-times” hypothesis argues that organisms deliberately re-allocate scarce resources away from reproduction to somatic maintenance. Indeed, DR commonly results in reduced fecundity, and this effect has been reported so often that reduced reproduction is sometimes included in the formal definition of DR^9^.

Nevertheless, the resource allocation hypothesis has been challenged in recent years by empirical studies showing that the relationship between lifespan extension and reduced reproduction under DR is not obligatory and can be uncoupled. Thus, Grandison et al.^9^ showed that addition of the dietary amino acid, methionine, restores fecundity of *Drosophila melanogaster* fruit flies under DR to normal levels without affecting their extended longevity. More generally, DR famously extends lifespan in sterile or non-reproducing animals^10–13^. Furthermore, experimental evolution studies suggest that longevity and reproduction do not necessarily co-evolve under DR^14,15^. These results suggest that animals under DR may not live longer because of resource re-allocation from reproduction to survival.

However, previous studies that investigate trade-offs under DR focused primarily on survival and fecundity of the parents and did not consider important traits such as offspring quality. Intergenerational trade-offs between fitness-related traits in parents and their offspring are well known from classic life-history theory^16,17^ but are rarely incorporated in the empirical analyses of the fitness consequences of DR. Increased investment into somatic maintenance can improve survival and even post-DR reproduction of organisms compared to non-starved controls. Indeed, there is some indication in the literature that DR can result in improved late-life reproduction^13^. Nevertheless, little is known about the quality of the resulting offspring, and life-history theory suggests that increased investment into current survival and reproduction can come at the cost of the propagule size^16,17^.

DR has been shown to robustly increase lifespan in *Caenorhabditis* nematodes using a broad variety of techniques^18^, including through temporary fasting by withholding food completely for a period of time^19,20^. Here we used the temporary fasting (TF) approach to investigate the effects of DR on survival, stress resistance and reproduction of females and their offspring in the nematode *Caenorhabditis remanei*. To test for the ‘missing costs’ of DR, we manipulated diet by either depriving females of a bacterial food source for six consecutive days or providing constant access to food. Following the treatment, we studied age-specific reproduction, survival and heat-shock resistance in mothers and their female offspring to investigate intergenerational effects of maternal temporary fasting. Building on the previous research and predictions derived from the life-history theory, we hypothesized that TF during early adulthood will positively affect fitness-related traits of ageing mothers, but there will be negative intergenerational effects that will reduce fitness of their daughters.

## Results

### Temporary fasting increases female fitness components

Females in the temporary fasting (TF) treatment produced a significantly higher number of offspring once returned to food and allowed mating (mean ± se; 463.45 ± 20.70) relative to females given complete access to food (330. 76 ± 22.73; Wald Z = 19.33, *P* < 0.005). There was also an interaction between age at last reproduction and diet treatment (Z = 3.12, *P* < 0.002) with TF females reproducing to a later age (14.71 ± 0.34 days) than *ad libitum* (AL) females (12.50 ± 0.34 days). Though TF females produce a higher median number of offspring per day, there was no statistical difference in daily reproductive output (Z = 0.632, *P* > 0.53, Fig. 1a). In line with fecundity, lifespan and heat stress resistance differed between treatments. Females that were deprived of bacteria had a significantly longer lifespan (median (95% CI), TF: 17 [15, 22] days; Coxph model, Wald’s Z= 2.497, p < 0.013) relative to females that had access to food (13.5 [13, 16] days; Fig. 2a). Similarly, females deprived of bacteria were more heat stress resistant relative to AL females (Wald’s Z=3.939, *P* < 0.001). This effect is dependent upon the length of exposure (Wald’s z= 2.192, *P* < 0.03), with a difference between TF and AL females up to the third hour of exposure (Fig. 2c).

**Figure 1.**
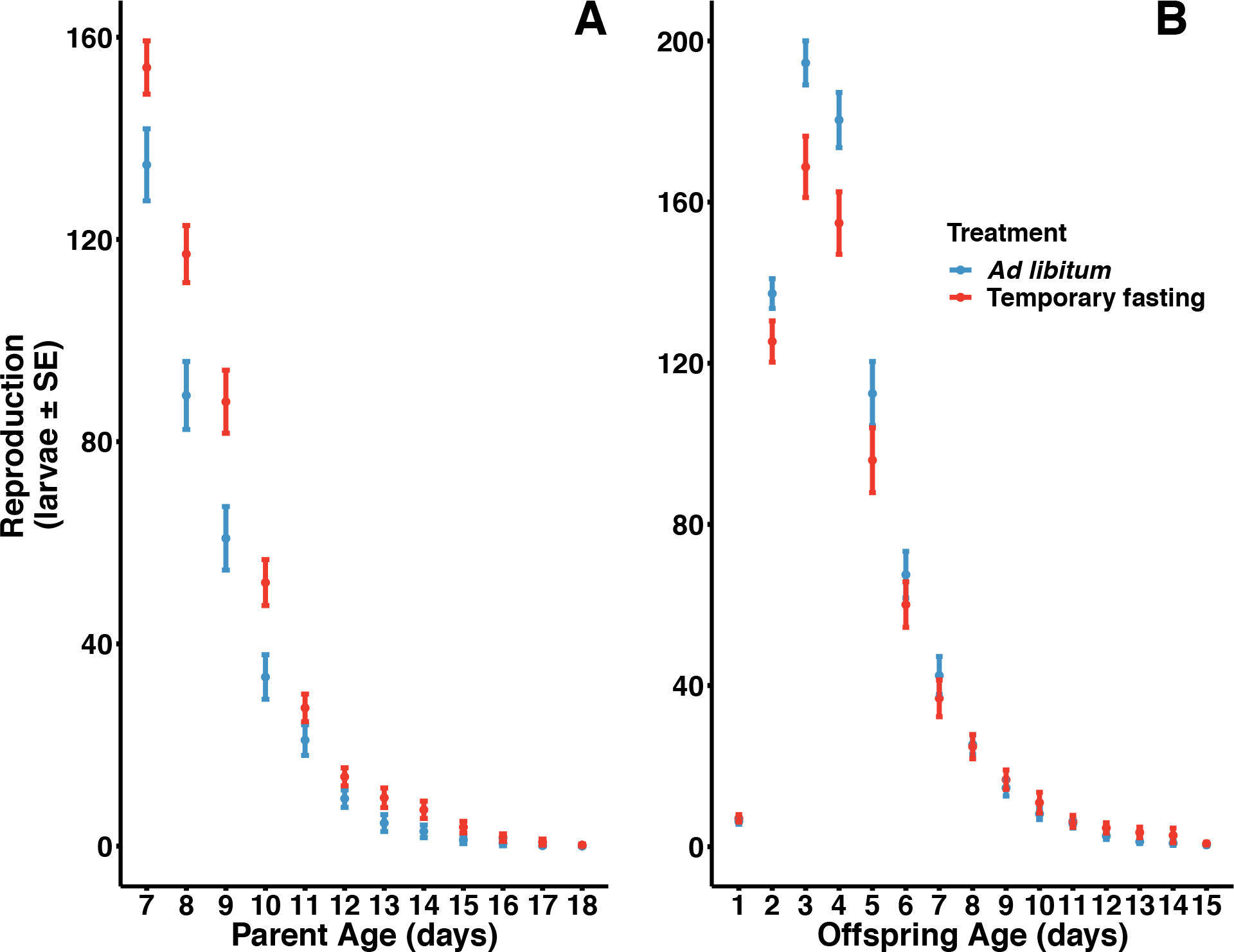
The influence of temporary fasting on fitness components of treatment females (left) and their offspring (right). The influence of BD on age-specific reproduction in treatment females (A) and their offspring (B).

**Figure 2.**
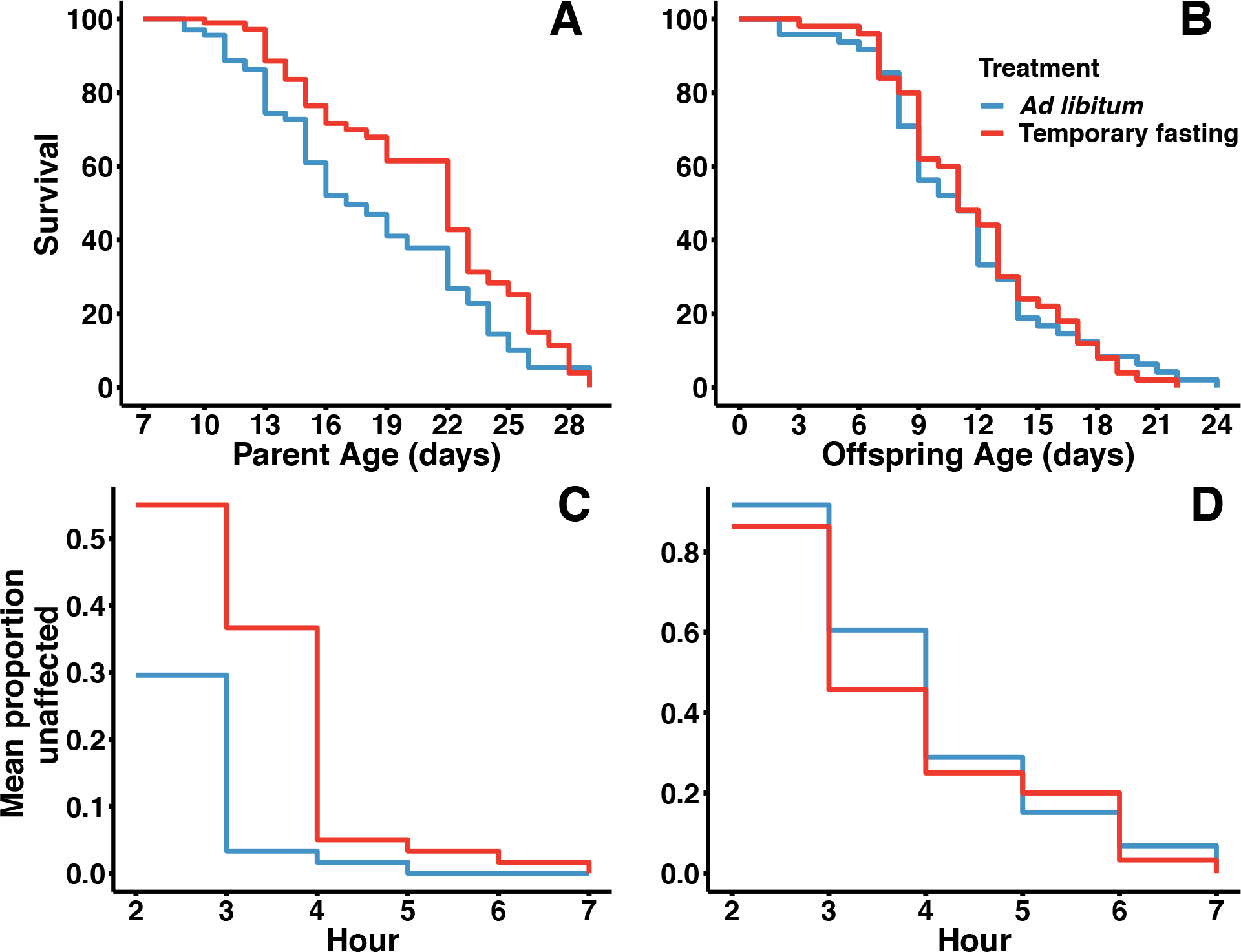
The impact of temporary fasting on female (A) and offspring (B) survival. The effect of BD on heat stress resistance in females (C) and offspring (C).

### Temporary fasting has negative or neutral effects on offspring

Offspring from mothers deprived of bacteria show reduced daily reproductive output during peak reproduction (D2 - D4) relative to offspring from mothers in the AL treatment (Fig. 1b). There was a significant quadratic effect of age in daily reproduction (Wald’s Z = 16.66, *P* < 0.001) and this quadratic effect interacted with diet treatment (Wald’s Z = 3.846, *P* < 0.001). In addition, total lifetime reproduction differed between treatments (Wald’s Z = 17.41, *P* < 0.0001). Offspring from AL mothers had a higher lifetime fecundity (751.67 ± 32.96) relative to offspring from TF mothers (690.90 ± 34.46). There was no difference in age at last reproduction (Wald’s Z = 3.868, *P* > 0.65). Maternal diet treatment did not influence offspring lifespan (Wald’s Z = 0.096, *P* = 0.924; median [95% CI], TF: 11 [9, 12]; AL: 11 [9, 13], Figure 2b). There was also no difference in heat stress resistance based on maternal diet (Wald’s Z = −0.892, *P* = 0.372, Figure 2d). In contrast, maternal diet has a significant effect on both development time and body size (Fig. 3a,b). Offspring from mothers in the TF treatment took significantly longer to develop (68.6 ± 0.22 hours) than offspring from mothers in the AL treatment (67.2 ± 0.22; t= −3.07, p = 0.002). Similarly, offspring from females in the temporary fasting experiment were significantly smaller (0.0293 ± 0.00022 mm^2^) than those with mothers in the *ad libitum* treatment (0.0302 ± 0.00025; t= 2.78, p < 0.0058).

**Figure 3.**
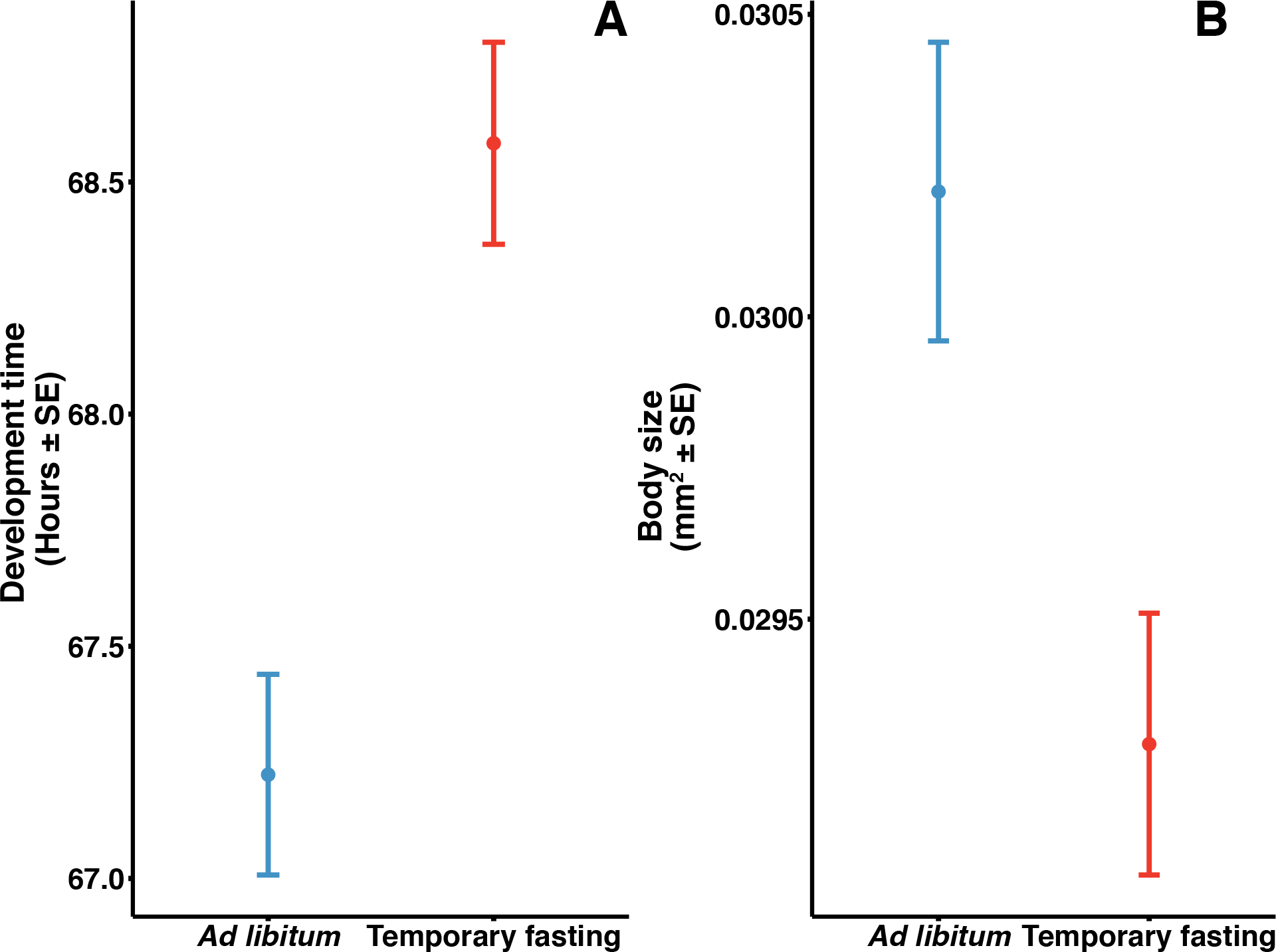
The impact of maternal temporary fasting treatment on offspring development time (a) and body size (b).

## Discussion

Temporary fasting had a strong positive effect across the three fitness related traits that we measured in females. Consistent with previous research in nematodes^18–20^ and other animals^6,21^, the dietary restriction by temporary fasting (DR by TF) treatment extended mean lifespan in females. Additionally, DR increased stress resistance as TF females also showed increased resistance to high temperature relative to control females, aligning with previously reported links between heat-shock resistance and lifespan in nematodes^22,23^. TF females also showed an increase in total lifetime fecundity and reproduced for longer than females that had constant access to food. The differences in reproduction and lifespan were not associated with a change in reproductive schedule as there was no difference in age-specific fecundity. At face value, these results align with the idea that lifespan can be extended through reduced food intake, without a corresponding cost to reproduction as has previously been suggested^21,24–27^.

Despite clear positive effects of DR on maternal survival in different environments and on lifetime reproductive success, none of these benefits were transferred to their female offspring. On the contrary, daughters of TF females show reduced initial reproduction (Figure 1b) and no increase in late-life reproductive output. As such, offspring from TF females show overall lower lifetime fecundity relative to offspring from females with constant access to food. Consistent with and compounding lower fecundity, offspring from TF mothers were smaller and took on average an hour and a half longer to develop to sexual maturity. We failed to detect any positive effects that might counter-balance these negative effects in offspring. There was neither a difference in survival in normal temperature nor a difference in heat-shock resistance between female offspring of TF and control mothers. Our results stand in contrast to a previous study in *C. elegans* that showed that DR offspring show increased larval resistance to starvation and produce more offspring relative to control worms^28^, suggesting that DR in mothers adaptively primes their offspring. We did not explicitly test for adaptive effects under offspring nutrient stress that mimic maternal food environments, though our results from heat-shock trials provide circumstantial evidence arguing against a generalized adaptive stress priming effect in *C. remanei*. More research is needed to determine if there are more specific priming effects under similar nutrients conditions as mothers.

The negative effects in offspring are a result of maternal dietary treatment, as offspring were reared in the same full, *ad libitum* (AL) diet. The effects on offspring can be explained by changes in maternal gamete provisioning^29,30^. There are well-known trends across taxa of both a trade-off between offspring number and size, as well as a positive correlation between gamete size and adult size^17,31^. Our results of increased fecundity of DR females and smaller adult size of their offspring are consistent with this explanation. This explanation is also congruent with the evidence from *C. elegans* of DR females showing lower fecundity, but larger investment in gamete size^28^. Different species can have different responses to DR in terms of their reproductive strategies; however, different DR regimes can also produce different types of response with respect to allocation decisions between investment in propagule number versus propagule size. Nevertheless, both scenarios highlight the importance of intergenerational trade-offs in the evolution of DR response. Our results show that dietary restriction can indeed improve fitness-related traits in ageing mothers but at the cost of reduced fitness-related traits in their daughters. Different forms of DR, including temporary fasting, as well as different DR mimetics, are increasingly used in humans both in clinical practice and in research. Future studies should incorporate intergenerational effects of DR across a broad range of taxa to improve our understanding of the potential “missing costs” of the beneficial DR effects.

## Methods

We used wild type *Caenorhabditis remanei* SP8 strain worms that were maintained for 50 generations under standard conditions of 20°C and 60% humidity and later stored in a −80°C freezer before use in experiments. Worms were thawed and maintained at 20°C for an additional 5 generations under standard lab conditions on NGM agar plates containing kanamycin and streptomycin and seeded with *E. coli* OP50-1 (pUC4K) bacteria as a food source. Plates were kept in a Panasonic MLR-352H Environmental Chamber under standard conditions. Worms were bleached twice, 4 and 8 days after thawing, to standardize for development time and to kill unwanted microbes.

Two days after the second bleaching, L4-female larvae were placed in groups of 10 on standard NGM agar plates with food and allowed to complete sexual maturation overnight. Early the following morning, adult virgin females were split between two treatments of either *ad libitum* access to food (AL) or without access to any food (temporary fasting, TF). Female worms were maintained for 6-days in their respective diet treatments. The worms were moved to fresh plates approximately every 48 hours and 24 hours before being placed in assays. We performed a series of assays to investigate the effect of diet treatment on an array of fitness components of treatment females and of their offspring.

### Round 1: Treatment female lifespan and reproduction

From each food treatment, 50 females were place individually on standard NGM plates (with antibiotics) seeded with standard food. Females were placed with 2 males (1-3 days post sexual maturity) derived from SP8-50 stock populations. New males of the same age were provided every 3rd day. To assess reproduction and lifespan, females and males were transferred to a new plate every 24h. The plate harbouring eggs from the previous 24-hour laying period were kept under standard conditions for an additional ~30 hours to allow offspring to hatch. After this period, larvae were euthanized in an environmental chamber at 39°C for 2 hours. Plates were then immediately placed in a refrigerator at 4°C until larval counts took place. Larval counts were performed within 24 hours after euthanization. After 13 days, reproduction had ceased, and females were transferred to new plates every third day, but still checked daily for lifespan scoring. Females were maintained as above until death. We recorded the date of death for each female. One female in the *ad libitum* treatment died from dehydration after climbing the plastic side of the petri dish and was excluded from the data.

### Heat stress resistance

To assess heat stress resistance, we placed 6 plates of 10 females from each treatment (12 plates total) in an environmental chamber at 37.5°C. Starting 2 hours after and every hour up to 7 hours after initial exposure, females were monitored for a knockdown response. At each time point, females were scored and placed in one of three categories following Herndon et al.^32^: (1) Freely moving or normal movement after being prodded gently by scorer; (2) Abnormal, non-sinusoidal movement or not freely moving (i.e. only anterior portion of body moving) after being prodded gently by scorer; and (3) not moving even after being stimulated by scorer, but with buccal pump moving or completely non-response. Females scored in categories 2 and 3 were combined later for analytical purposes. After the 7-hour heat shock period, plates were stored at 20°C and females reassessed using the same categories for recovery ~ 21 hours after final 7^th^ hour scoring.

### Effects on offspring reproduction and lifespan

We collected 1 L4 stage female larvae from the first 24-hour laying period from each female in both treatments. Reproduction and lifespan assays followed the same procedure as the mothers above, but female offspring were immediately placed into reproduction assays after collection to ensure reproduction commenced as soon as they reached sexual maturity. We recorded daily measures of larval output and the date of death for each female offspring.

### Round 2

The second round was aimed to collect additional offspring traits. In the second round, after 6 days of diet exposure, 10 treatment females were placed on seeded standard NGM plates that had 10, 2-day (post-sexual maturity) adult males such that there was 1:1 sex ratio. Females were kept on these plates for 2 hours to allow for feeding, mating, and for egg laying to commence. Any eggs laid during this period were discarded. Afterwards, females and males were transferred to a new seeded NA plate and allowed to lay eggs for 1-hour. This procedure was performed a second time on a new empty, seeded plate. The eggs from the two separate hour-long periods were used to estimate offspring development time and body size at maturity (plate 1), as well as to assess heat stress resistance (plate 2). Treatment females and males were removed and discarded after the two laying periods.

### Development time and size at maturity

The offspring from the first plate were allowed to mature on the laying plates. On the 3^rd^ day post egg laying, we monitored plates hourly for sexually mature females. Females were removed from the plates either at the time the vulva showed full maturation (i.e. when the two lips of the vulva were visible as bumps) or if they appeared to have mated with males present on the plate. These two events typically occur in close proximity. We recorded the (whole) hour at which each female was collected as sexually mature and subtracted it from the time at which mothers were removed from the lay plate to estimate development time. Plates were monitored until there were no females remaining. Immediately after females were removed from the plates (i.e. at sexual maturity), we photographed them using a Lumenera Infinity2-5C digital microscope camera mounted on a Leica M165C stereomicroscope. These photos were then used to measure body area as an estimate of body size. We followed the methods of^33^. Briefly, photos were imported to ImageJ (v. 1.50i; https://imagej.nih.gov/ij/). An initial image with a stage micrometer was used to calibrate the length of a pixel at the settings used to take photos of offspring. Images of each offspring were imported and converted to gray scale. The threshold function was used to provide a physical outline of the worm’s body. Any area not highlighted within the body of the worm after this procedure was then colored in manually. The total area was then estimated using the measure function in ImageJ.

### Heat stress resistance

Virgin females were collected from the second plate on the 3^rd^ day post-laying (as above). We isolated two plates of 10 females from each of the 3 laying plates in each treatment (for a total of 12 plates). On day 4 post-sexual maturity, females were moved to new plates and placed immediately in to heat stress treatment. We followed the same protocols that were used for the parental generation in experiment 1 to assess the response of offspring.

### Statistical analysis

Data were analysed using *R 3.4.0* statistical software^34^. Age-specific and lifetime reproductive output were analysed using a multilevel mixed model in the *lme4* package^35^ with a Poisson error distribution, log-link function. Both age and age^2^ were included in the model to asses linear and quadratic effects of this factor (see Supplemental Material). Individual (to account for repeated measures) and individual record value (to account for any overdispersion^36^) were included as random effects for age-specific analysis; only line was used as a random effect in reproduction analyses. Heat stress resistance was also analysed using a multilevel model but using a binomial error distribution and logit link function. We included a random effect of ‘plate’ to account for variation of plates on which females were treated. Development time and body size were analysed using a general linear model. Lifespan was analysed using a Cox Proportional hazards model in the survival package^37^.

## Supporting information

Supplemental Material Statistical Report

## Acknowledgements

This research was funded by the European Research Council AgingSexDiff and GermlineAgeingSoma and Swedish Research Council to AAM. The authors would like to thank Hanne Carlsson for her assistance.

## Author contributions

A.M. conceived of the project and contributed to the manuscript. A.M. & B.M designed the experiment. M.L. executed the experiment and contributed to the manuscript. B.M. executed the experiments, analysed the data, and wrote the manuscript.

## Additional information

Supplementary information is available for this paper.

Correspondence for this paper can be sent to B.M and request for material to A.M.

