## Supplemental Material Statistical Report for "Dietary restriction improves fitness of ageing parents but reduces fitness of their offspring"

### Effects of Temporary Fasting on female fitness and on offspring in *Caenorhabditis remanei*

*Brian Mautz*

*2017-05-15*

#### Contents

#**Effects on focal females (F0)** Below is an analysis of the Temporary Fasting experiments.

##**F0 AGE-SPECIFIC REPRODUCTION** ###*Analysis of age-specific reproduction* *upload data, upload packages, inspect data (code hidden)

###Descriptive statistics and initial plots

```
dtworm[treatment == 'AL', .(meanAL_larv = mean(larvae), sdAL_larv= sd(larvae), iqrAL_larv= IQR(larvae))
```

```
##     plate  meanAL_larv  sdAL_larv iqrAL_larv
##  1:     1 134.76000000 50.3512398      71.00
##  2:     2  89.14000000 47.5321631      67.75
##  3:     3  60.88636364 41.6805213      64.75
##  4:     4  33.45238095 28.5616171      39.00
##  5:     5  20.97142857 17.9320379      26.00
##  6:     6   9.39393939  9.9528243      16.00
##  7:     7   4.56000000  8.3570330       5.00
##  8:     8   2.91666667  6.1071353       4.25
##  9:     9   1.22222222  3.0784938       0.75
## 10:    10   0.57142857  1.8693596       0.00
## 11:    11   0.07692308  0.2773501       0.00
## 12:    12   0.00000000  0.0000000       0.00
## 13:    13   0.00000000  0.0000000       0.00
## 14:    14   0.00000000  0.0000000       0.00
## 15:    15   0.00000000  0.0000000       0.00
## 16:    16   0.00000000  0.0000000       0.00
## 17:    17   0.00000000  0.0000000       0.00
## 18:    18   0.00000000  0.0000000       0.00
## 19:    19   0.00000000  0.0000000       0.00
## 20:    20   0.00000000         NA       0.00
## 21:    21   0.00000000         NA       0.00
## 22:    22   0.00000000         NA       0.00
##     plate  meanAL_larv  sdAL_larv iqrAL_larv
```

```
dtworm[treatment == 'TF', .(meanBD_larv = mean(larvae), sdBD_larv= sd(larvae), iqrBD_larv= IQR(larvae))
```

```
##     plate meanBD_larv  sdBD_larv iqrBD_larv
##  1:     1 154.0000000 36.9126221      39.00
##  2:     2 117.1020408 39.5365047      45.00
##  3:     3  87.8979592 43.6468999      62.00
##  4:     4  52.1276596 31.1731609      46.50
##  5:     5  27.3404255 18.7248691      32.50
##  6:     6  13.7111111 11.9917059      16.00
##  7:     7   9.5675676 11.8169853      15.00
##  8:     8   7.1818182  9.8407779       7.00
```

```
##  9:      9  3.7500000  6.0530066       6.25
## 10:     10  1.6800000  3.6823905       1.00
## 11:     11  0.7916667  2.9337491       0.00
## 12:     12  0.2173913  0.8504823       0.00
## 13:     13  0.1500000  0.4893605       0.00
## 14:     14  0.0000000  0.0000000       0.00
## 15:     15  0.0500000  0.2236068       0.00
## 16:     16  0.0000000  0.0000000       0.00
## 17:     17  0.0000000  0.0000000       0.00
## 18:     18  0.0000000  0.0000000       0.00
## 19:     19  0.0000000  0.0000000       0.00
## 20:     20  0.0000000  0.0000000       0.00
## 21:     21  0.0000000  0.0000000       0.00
## 22:     22  0.0000000         NA       0.00
##      plate meanBD_larv  sdBD_larv iqrBD_larv
```

```
interact<-interaction(dtworm$treatment, dtworm$plate, sep=":")
ggplot(data=dtworm, aes(x=interact, y=larvae)) +
  geom_boxplot(aes(fill=dtworm$treat), width = 0.6) +
  theme_bw() +
  theme(axis.text.x=element_text(angle=90, hjust = 1), panel.grid.major = element_blank(), panel.grid.mi
        panel.background = element_blank(), axis.line = element_line(colour = "black")) +
  ggtitle("Lifetime offspring production by day in treatment worms")
```

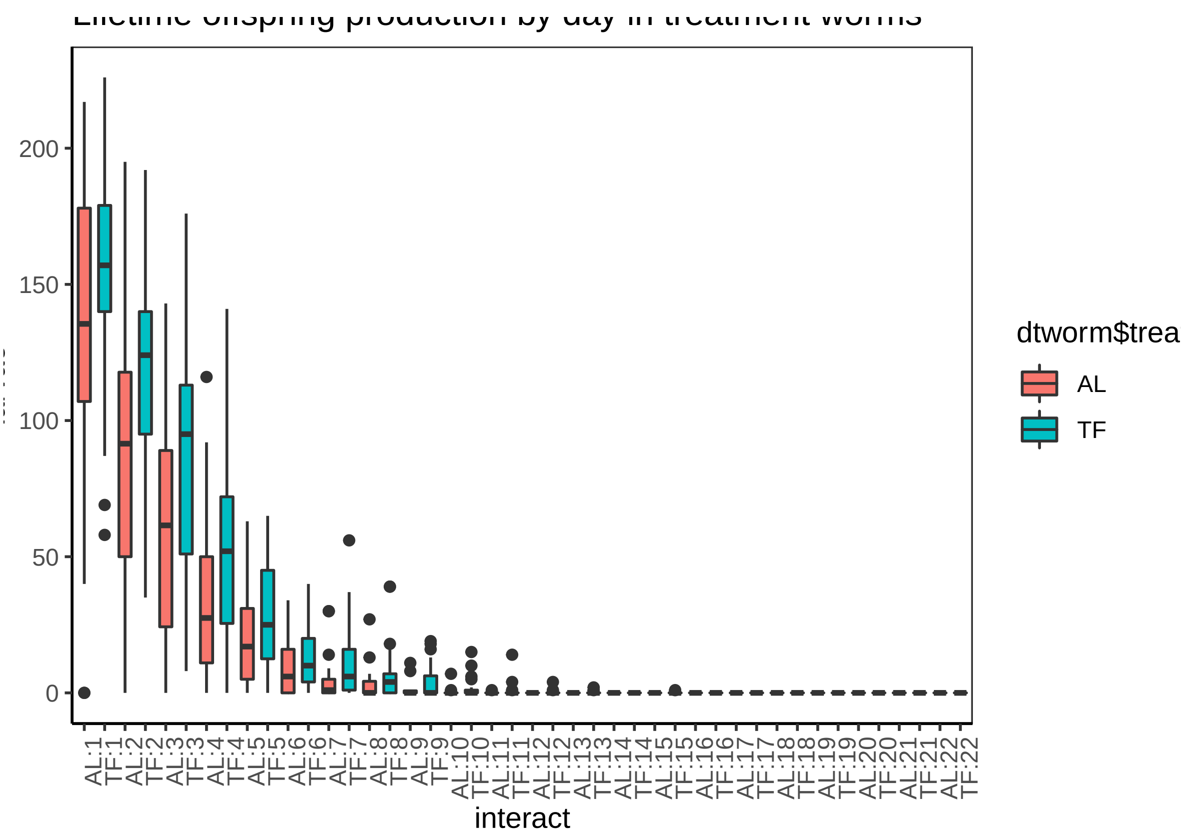


#####Initial model doesn't converge

```
dtworm <- dtworm[dtworm$plate < 16, ]
```

```r
m1<-glmer(dtworm$larvae~ (dtworm$pltstd1 + dtworm$pltsqstd)*dtworm$treatment + dtworm$alr2std*dtworm$tre
```

```
## Warning in checkConv(attr(opt, "derivs"), opt$par, ctrl =
## control$checkConv, : Model failed to converge with max|grad| = 0.144813
## (tol = 0.001, component 1)
```

###### Understanding convergence issue

Model converges with updating m1 'theta' and 'fixef' variables, increasing iterations, and changing optimizer function

###Final model that converges. There was a significant difference in the lifetime number of larvae produced between treatments. Females in the Temporary Fasting treatment produced a significantly higher number of offspring relative to female given access to food (Wald Z= 2.81, $p < 0.005$, Figure 1). We detected a significant quadratic effect of reproduction (Z= 2.91, $p < 0.004$), but the shape and rate of reproductive decline did not differ between treatments (Z= 0.632, $p > 0.53$). There was also an interaction between age and last reproduction and diet treatment (Z=3.12, $p < 0.002$) with the average age at last reproduction lower in bacterially deprived females.

```r
###########################Try a different optimizer###################
m3<-update(m2, start=ss, control=glmerControl(optimizer="bobyqa",optCtrl=list(maxfun=2e5)))
    # model still does not converge, 11 negative eigenvalues
summary(m3)
```

```
## Generalized linear mixed model fit by maximum likelihood (Laplace
##   Approximation) [glmerMod]
##  Family: poisson  ( log )
## Formula:
## dtworm$larvae ~ (dtworm$pltstd1 + dtworm$pltsqstd) * dtworm$treatment +
##     dtworm$alr2std * dtworm$treatment + (1 | dtworm$individual) +
##     (1 | dtworm$line)
##    Data: dtworm
## Control:
## glmerControl(optimizer = "bobyqa", optCtrl = list(maxfun = 2e+05))
##
##      AIC      BIC   logLik deviance df.resid
##   6412.7   6460.8  -3196.4   6392.7      894
##
## Scaled residuals:
##     Min      1Q  Median      3Q     Max
## -1.8664 -0.2729 -0.0328  0.0715  5.9666
##
## Random effects:
##  Groups            Name        Variance Std.Dev.
##  dtworm$line       (Intercept) 0.8135   0.9019
##  dtworm$individual (Intercept) 0.1193   0.3454
## Number of obs: 904, groups:  dtworm$line, 904; dtworm$individual, 50
##
## Fixed effects:
##                                    Estimate Std. Error z value Pr(>|z|)
## (Intercept)                          0.4822     0.2385   2.021  0.04324
## dtworm$pltstd1                      -2.4490     0.4198  -5.833 5.43e-09
## dtworm$pltsqstd                     -2.3497     0.8105  -2.899  0.00374
## dtworm$treatmentTF                   0.7274     0.2578   2.822  0.00477
## dtworm$alr2std                       0.6844     0.0788   8.686  < 2e-16
```

```
## dtworm$pltstd1:dtworm$treatmentTF    0.3670     0.5104   0.719  0.47208
## dtworm$pltsqstd:dtworm$treatmentTF   0.6082     0.9354   0.650  0.51556
## dtworm$treatmentTF:dtworm$alr2std   -0.3134     0.1005  -3.119  0.00181
##
## (Intercept)                        *
## dtworm$pltstd1                     ***
## dtworm$pltsqstd                    **
## dtworm$treatmentTF                 **
## dtworm$alr2std                     ***
## dtworm$pltstd1:dtworm$treatmentTF
## dtworm$pltsqstd:dtworm$treatmentTF
## dtworm$treatmentTF:dtworm$alr2std  **
## ---
## Signif. codes:  0 '***' 0.001 '**' 0.01 '*' 0.05 '.' 0.1 ' ' 1
##
## Correlation of Fixed Effects:
##             (Intr) dtwr$1 dtwrm$ dtw$TF dtwr$2 d$1:$T d$:$TF
## dtwrm$plts1 -0.743
## dtwrm$pltsq  0.879 -0.952
## dtwrm$trtTF -0.874  0.687 -0.808
## dtwrm$lr2st  0.068 -0.095 -0.003 -0.075
## dtwrm$1:$TF  0.610 -0.821  0.782 -0.728  0.073
## dtwrm$p:$TF -0.756  0.824 -0.864  0.877  0.003 -0.951
## dtwrm$TF:$2 -0.042  0.065  0.006 -0.026 -0.715 -0.054 -0.028
```

##*F0 net reproductive fitness analysis* ####*Subset age-specific data

####Initial table and plot

```
#descriptive stat and boxplot of LRS by treatment
lfspn[, .(mean_LRS = mean(ttllrv), median_ttllrv = median(ttllrv), seAL_ttlrv = sd(ttllrv)/sqrt(length(
```

```
##    treatment mean_LRS median_ttllrv seAL_ttlrv iqr_LRS
## 1:        AL  330.760           329   22.72904   236.5
## 2:        TF  463.449           464   20.70400   217.0
```

```
#descriptive stats for age at last reproduction
lfspn[, .(mean_LRS = mean(alr), seAL_lfspn = sd(alr)/sqrt(length(individual)), iqr_LRS = IQR(alr)), by=
```

```
##    treatment mean_LRS seAL_lfspn iqr_LRS
## 1:        AL 12.50000  0.3431546       3
## 2:        TF 14.71429  0.3413160       3
```

```
ggplot(lfspn, aes(treatment, ttllrv, color=treatment))+
  geom_boxplot(width = 0.6) +
  theme_bw() +
  theme(axis.text.x=element_text(angle=90, hjust = 1), panel.grid.major = element_blank(),        panel
        panel.background = element_blank(), axis.line = element_line(colour = "black")) +
  ggtitle("Lifetime offspring production in treatment worms")
```

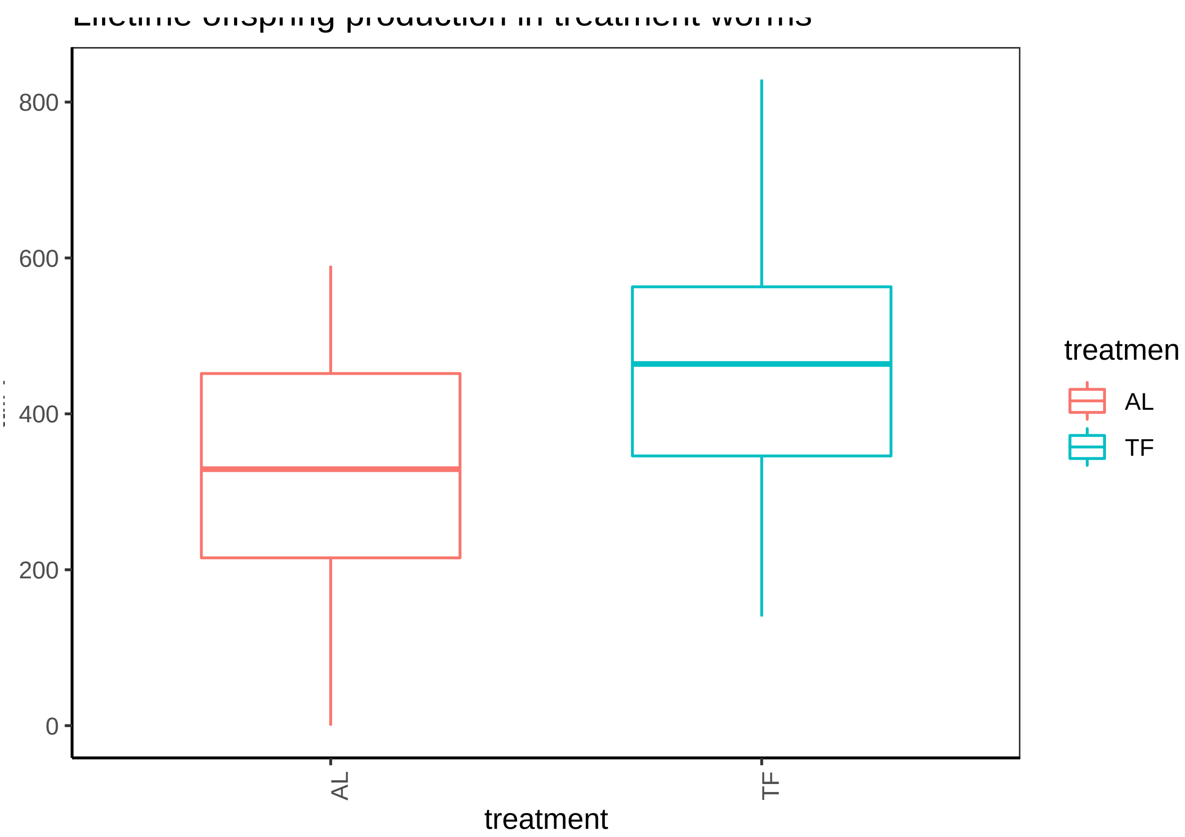


###Statistical model

```
m2 <- glmer(ttllrv ~ treatment + (1|lifespan), data=lfspn, family='poisson')
summary(m2)
```

```
## Generalized linear mixed model fit by maximum likelihood (Laplace
##   Approximation) [glmerMod]
##  Family: poisson  ( log )
## Formula: ttllrv ~ treatment + (1 | lifespan)
##    Data: lfspn
##
##      AIC      BIC   logLik deviance df.resid
##   5501.7   5509.5  -2747.9   5495.7       96
##
## Scaled residuals:
##      Min       1Q   Median       3Q      Max
## -19.4345  -4.1191   0.0216   3.7098  14.1896
##
## Random effects:
##  Groups   Name        Variance Std.Dev.
##  lifespan (Intercept) 0.09175  0.3029
## Number of obs: 99, groups:  lifespan, 20
##
## Fixed effects:
##             Estimate Std. Error z value Pr(>|z|)
## (Intercept)  5.85732    0.06834   85.70   <2e-16 ***
## treatmentTF  0.22244    0.01151   19.33   <2e-16 ***
## ---
```

```
## Signif. codes:  0 '***' 0.001 '**' 0.01 '*' 0.05 '.' 0.1 ' ' 1
##
## Correlation of Fixed Effects:
##              (Intr)
## treatmentTF -0.094
```

##**F0 LIFESPAN** ####Descriptive statistic and initial plot

```
##    treatment mean_lfspn median_lfspln sd_lfspn iqr_lfspn
## 1:        AL   15.14000          13.5 5.134954      6.75
## 2:        TF   18.44898          17.0 5.412413      9.00
```

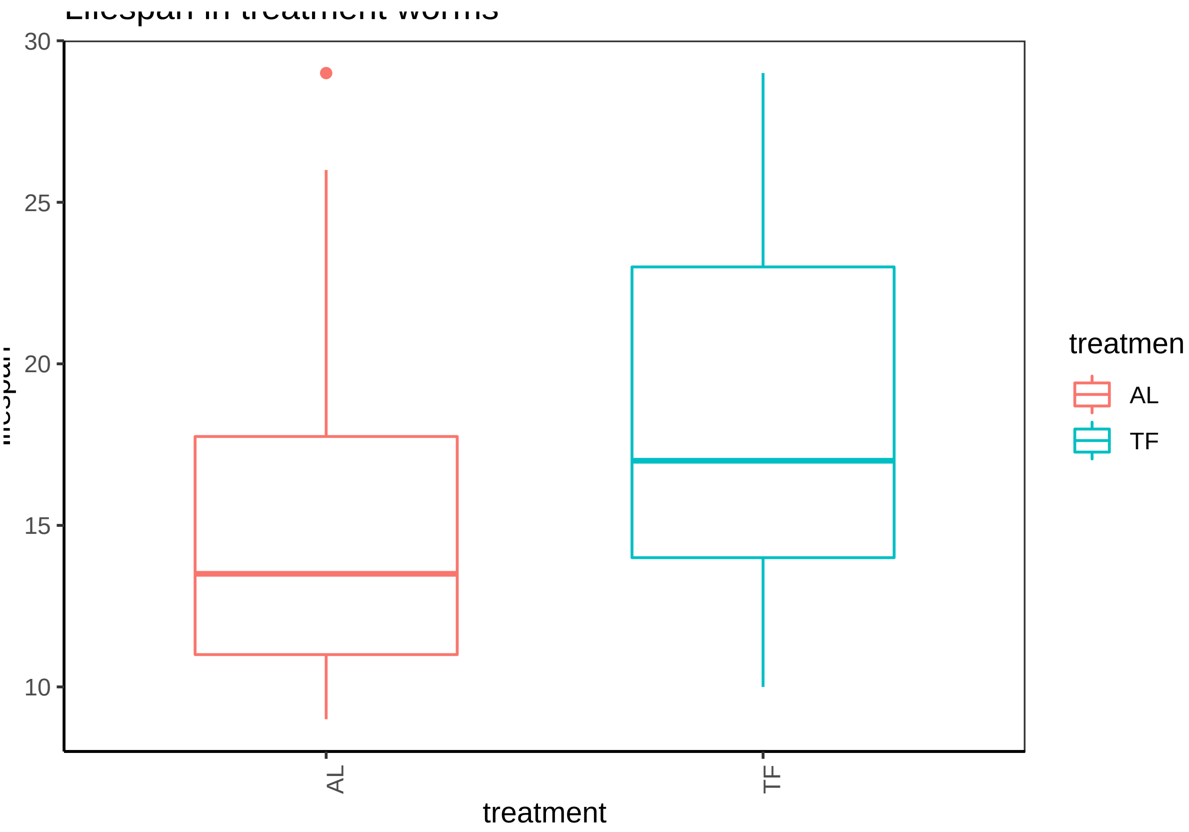

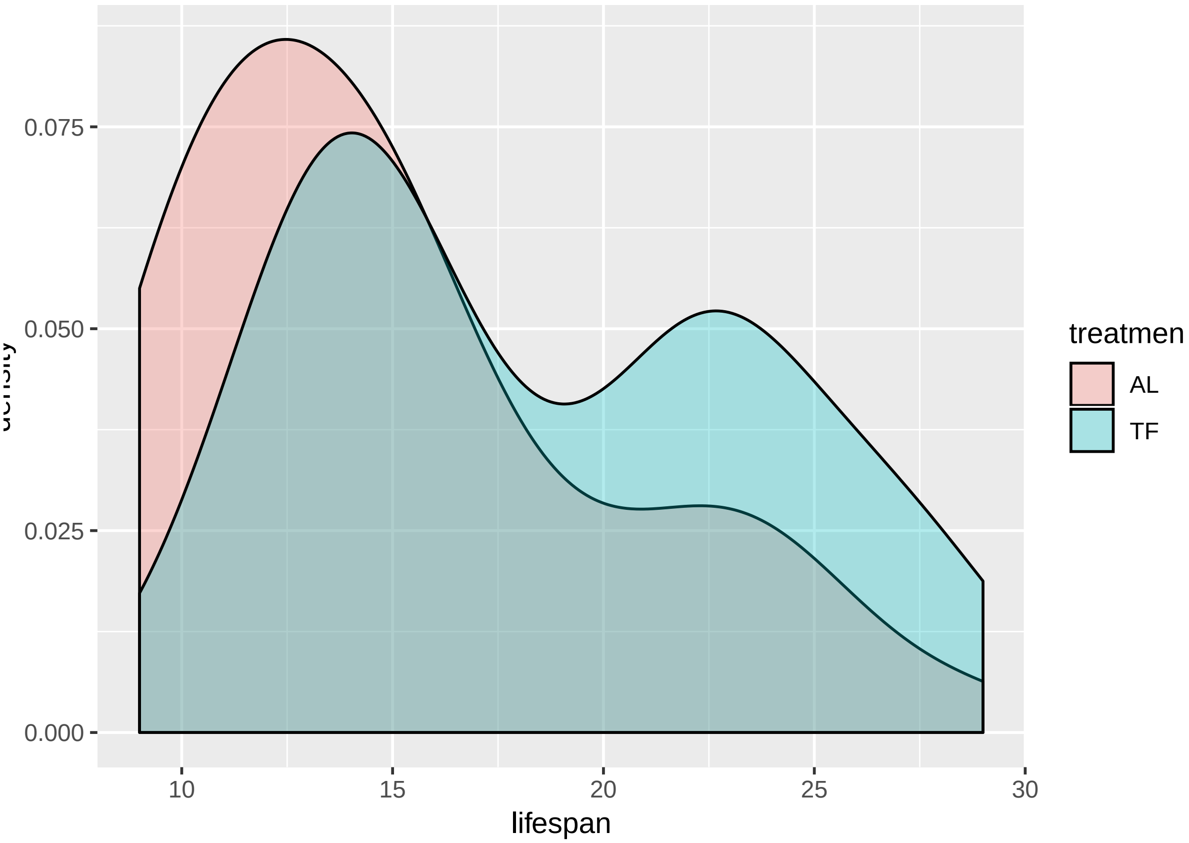


###Analysis Females that were deprived of bacteria had a significantly longer lifespan (median ± sd; ; BD: 17 ± 5.4 days; Coxph model, Wald's Z= 2.497, p< 0.013) relative to females that had access to food (AL: 13.5 ± 5.1 days; Figure 2).

```
#ANALYSIS USING COXPH IN SURVIVAL package
m1 <- coxph(Surv(lifespan,Event)~treatment ,data=lfspn)
summary(m1)
```

```
## Call:
## coxph(formula = Surv(lifespan, Event) ~ treatment, data = lfspn)
##
##   n= 99, number of events= 99
##
##                 coef exp(coef) se(coef)      z Pr(>|z|)
## treatmentTF -0.5079    0.6018   0.2034 -2.497   0.0125 *
## ---
## Signif. codes:  0 '***' 0.001 '**' 0.01 '*' 0.05 '.' 0.1 ' ' 1
##
##             exp(coef) exp(-coef) lower .95 upper .95
## treatmentTF    0.6018      1.662    0.4039    0.8965
##
## Concordance= 0.597  (se = 0.029 )
## Likelihood ratio test= 6.19  on 1 df,   p=0.01
## Wald test            = 6.24  on 1 df,   p=0.01
## Score (logrank) test = 6.36  on 1 df,   p=0.01
```

```
#FITTING BASIC MODEL USING SURVFIT to plot
survival.female <- survfit(Surv(lifespan,Event)~treatment, data=lfspn)
```

```r
summary(survival.female)
```

```
## Call: survfit(formula = Surv(lifespan, Event) ~ treatment, data = lfspn)
##
##                 treatment=AL
##  time n.risk n.event survival std.err lower 95% CI upper 95% CI
##     9     50       6     0.88  0.0460      0.79438        0.975
##    10     44       2     0.84  0.0518      0.74429        0.948
##    11     42       7     0.70  0.0648      0.58384        0.839
##    12     35       2     0.66  0.0670      0.54093        0.805
##    13     33       8     0.50  0.0707      0.37896        0.660
##    14     25       1     0.48  0.0707      0.35971        0.641
##    15     24       6     0.36  0.0679      0.24877        0.521
##    16     18       4     0.28  0.0635      0.17952        0.437
##    17     14       1     0.26  0.0620      0.16289        0.415
##    18     13       1     0.24  0.0604      0.14655        0.393
##    19     12       2     0.20  0.0566      0.11489        0.348
##    20     10       1     0.18  0.0543      0.09962        0.325
##    22      9       3     0.12  0.0460      0.05665        0.254
##    23      6       1     0.10  0.0424      0.04354        0.230
##    24      5       2     0.06  0.0336      0.02003        0.180
##    25      3       1     0.04  0.0277      0.01029        0.156
##    26      2       1     0.02  0.0198      0.00287        0.139
##    29      1       1     0.00     NaN           NA           NA
##
##                 treatment=TF
##  time n.risk n.event survival std.err lower 95% CI upper 95% CI
##    10     49       2   0.9592  0.0283      0.90535        1.000
##    12     47       2   0.9184  0.0391      0.84482        0.998
##    13     45       8   0.7551  0.0614      0.64381        0.886
##    14     37       4   0.6735  0.0670      0.55417        0.818
##    15     33       5   0.5714  0.0707      0.44839        0.728
##    16     28       3   0.5102  0.0714      0.38779        0.671
##    17     25       1   0.4898  0.0714      0.36805        0.652
##    18     24       1   0.4694  0.0713      0.34853        0.632
##    19     23       3   0.4082  0.0702      0.29135        0.572
##    22     20       7   0.2653  0.0631      0.16649        0.423
##    23     13       4   0.1837  0.0553      0.10179        0.331
##    24      9       1   0.1633  0.0528      0.08662        0.308
##    25      8       1   0.1429  0.0500      0.07195        0.284
##    26      7       3   0.0816  0.0391      0.03192        0.209
##    27      4       1   0.0612  0.0342      0.02045        0.183
##    28      3       2   0.0204  0.0202      0.00293        0.142
##    29      1       1   0.0000     NaN           NA           NA
```

```r
survival.female
```

```
## Call: survfit(formula = Surv(lifespan, Event) ~ treatment, data = lfspn)
##
##                n events median 0.95LCL 0.95UCL
## treatment=AL 50     50   13.5      13      16
## treatment=TF 49     49   17.0      15      22
```

```r
plot(survival.female,lty=c(1,1),col=c('#F8766D','#00BFC4'),xlab="Adult age (days)",ylab = "Survival prol
legend("topright",c("Ad libitum","Temporary Fasting"), bty="n",lty=c(1,1),col=c('#F8766D','#00BFC4'))
```

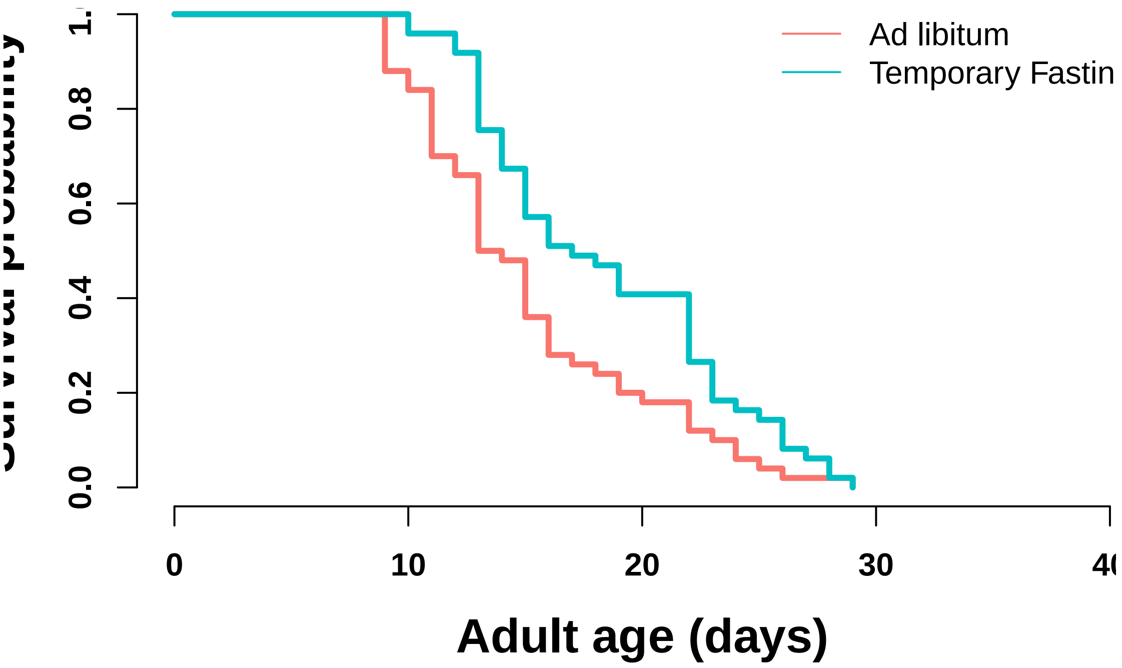


```
#check assumption of the ph model
temp <- cox.zph(m1)
m1
```

```
## Call:
## coxph(formula = Surv(lifespan, Event) ~ treatment, data = lfspn)
##
##                 coef exp(coef) se(coef)      z      p
## treatmentTF -0.5079    0.6018   0.2034 -2.497 0.0125
##
## Likelihood ratio test=6.19  on 1 df, p=0.01287
## n= 99, number of events= 99
```

##**F0 HEAT SHOCK** Data is analyzed by comparing individuals unaffected (response A, only) to those that appear affected (response B + respone C). These responses were bound (using cbind) and used as the response in a glmm with plate as a random factor and with a binomial error structure.

####Statistical analysis

There is signficant difference in the knockdown response of Temporary Fasting females relative to ad libitum females. Females deprived of bacteria are more heat stress resistant relative to ad lib females (Wald's Z=4.004, $p < 0.001$). This effect also is dependent upon the length of exposure (Wald's z= 2.197, $p < 0.03$). [NB: it doesn't show it here, but it appears that the significant difference in proportion is in the second check (i.e. hour 3)].

```
m1 <- glmer(response ~ treatment*hour + (1|plate), data=dt_F0hs, family="binomial")
summary(m1)
```

```
## Generalized linear mixed model fit by maximum likelihood (Laplace
##   Approximation) [glmerMod]
##  Family: binomial  ( logit )
## Formula: response ~ treatment * hour + (1 | plate)
##    Data: dt_F0hs
##
##      AIC      BIC   logLik deviance df.resid
```

```
##    460.1    472.3   -225.1    450.1       79
##
## Scaled residuals:
##    Min     1Q Median     3Q    Max
## -1.881 -1.352 -1.054  1.653  5.330
##
## Random effects:
##  Groups Name        Variance Std.Dev.
##  plate  (Intercept) 0.002656 0.05154
## Number of obs: 84, groups:  plate, 6
##
## Fixed effects:
##                  Estimate Std. Error z value Pr(>|z|)
## (Intercept)      -2.78115    0.38949  -7.140 9.30e-13 ***
## treatmenttf       1.81681    0.46124   3.939 8.18e-05 ***
## hour              0.18122    0.07780   2.329   0.0198 *
## treatmenttf:hour -0.21044    0.09602  -2.192   0.0284 *
## ---
## Signif. codes:  0 '***' 0.001 '**' 0.01 '*' 0.05 '.' 0.1 ' ' 1
##
## Correlation of Fixed Effects:
##             (Intr) trtmnt hour
## treatmenttf -0.841
## hour        -0.919  0.776
## trtmnttf:hr  0.744 -0.910 -0.810
```

#**Effects of maternal treatment on offspring (F1)**

##**F1 AGE-SPECIFIC REPRODUCTION**

```
##     plate meanAL_larv  sdAL_larv iqrAL_larv
##  1:     1   6.2916667  4.8770340       6.25
##  2:     2 137.2978723 25.3539422      32.00
##  3:     3 194.5434783 37.1350128      28.50
##  4:     4 180.3695652 46.4310043      43.25
##  5:     5 112.4782609 54.0114141      78.50
##  6:     6  67.5111111 38.9140207      47.00
##  7:     7  42.5000000 31.2264795      33.75
##  8:     8  25.3658537 15.5848582      20.00
##  9:     9  14.6176471 11.8679980      21.50
## 10:    10   8.1481481  7.3469183       9.50
## 11:    11   5.7200000  5.3969127       5.00
## 12:    12   2.6521739  4.0182390       4.00
## 13:    13   1.2500000  1.8439089       2.25
## 14:    14   0.8571429  1.9158104       0.75
## 15:    15   0.4444444  0.8819171       0.00
## 16:    16   0.0000000  0.0000000       0.00
## 17:    17   0.0000000  0.0000000       0.00
## 18:    18   0.0000000  0.0000000       0.00
## 19:    19   0.0000000  0.0000000       0.00
## 20:    20   0.0000000  0.0000000       0.00
## 21:    21   0.0000000  0.0000000       0.00
## 22:    22   0.0000000  0.0000000       0.00
## 23:    23   0.0000000         NA       0.00
##     plate meanAL_larv  sdAL_larv iqrAL_larv
```

```
##     plate  meanBD_larv  sdBD_larv iqrBD_larv
##  1:     1   7.04000000  6.5057588       7.00
##  2:     2 125.40000000 36.1775666      35.00
##  3:     3 168.74000000 53.8345197      48.00
##  4:     4 154.79591837 54.3069899      59.00
##  5:     5  95.93877551 56.3017940      77.00
##  6:     6  60.10204082 39.5791638      48.00
##  7:     7  36.83333333 31.5879948      22.50
##  8:     8  24.83333333 19.4797164      18.75
##  9:     9  16.65000000 15.1920188      12.50
## 10:    10  10.93548387 14.3083553      11.00
## 11:    11   6.30000000  8.1161822       6.75
## 12:    12   4.66666667  6.2670058       5.25
## 13:    13   3.54545455  6.1390807       3.75
## 14:    14   2.81250000  7.2038763       1.00
## 15:    15   0.76923077  1.2351684       1.00
## 16:    16   0.09090909  0.3015113       0.00
## 17:    17   0.33333333  1.0000000       0.00
## 18:    18   0.00000000  0.0000000       0.00
## 19:    19   0.25000000  0.5000000       0.25
## 20:    20   0.00000000  0.0000000       0.00
## 21:    21   0.00000000         NA       0.00
## 22:    22   0.00000000         NA       0.00
##     plate  meanBD_larv  sdBD_larv iqrBD_larv
```

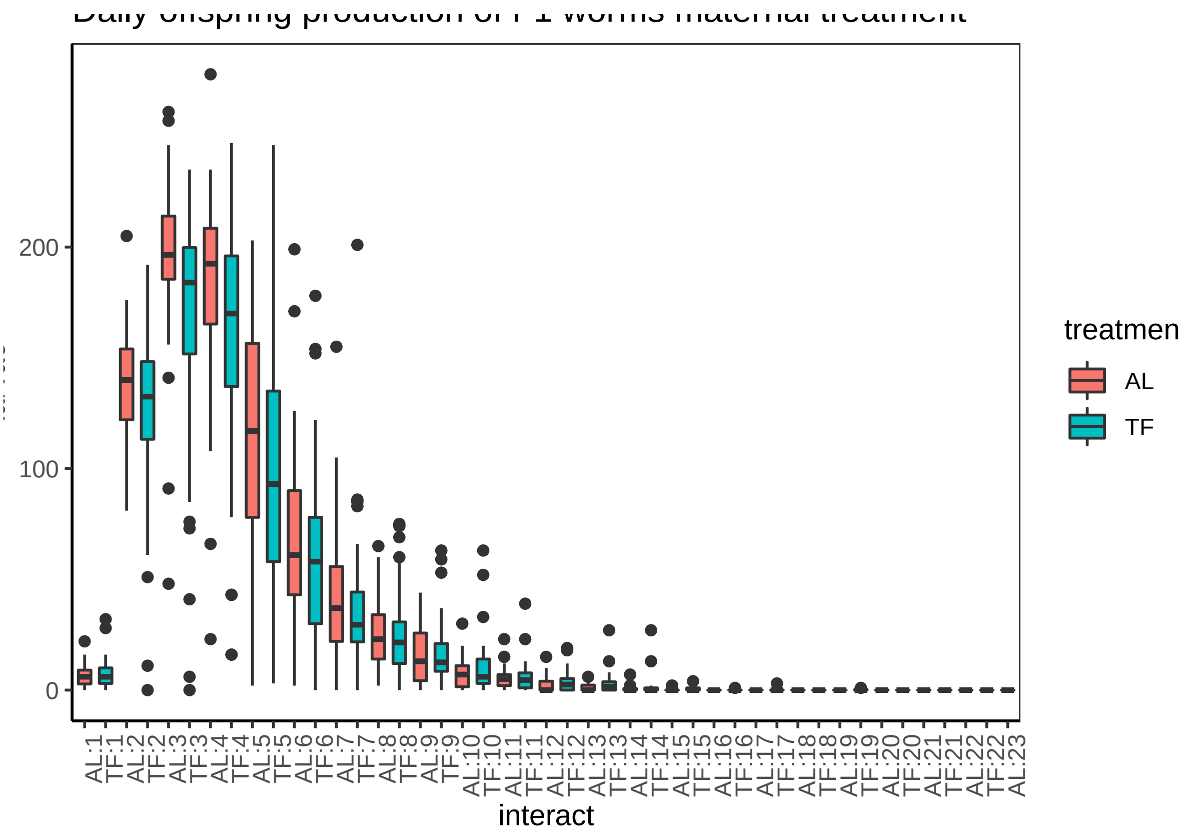


###*Analysis age-specific reproduction* Initial model fails to converge

```r
m1<-glmer(larvae~ (pltstd1 + pltsqstd)*treatment + alrstd*treatment + (1|individual) + (1|line), family=
```

```
## Warning in checkConv(attr(opt, "derivs"), opt$par, ctrl =
## control$checkConv, : Model failed to converge with max|grad| = 0.39747 (tol
## = 0.001, component 1)
```

###*Fixed convergence warning following https://rstudio-pubs-static.s3.amazonaws.com/33653_57fc7b8e5d484c909b615d8633c01d51.html#

###Final Model as above but increase number of iterations and changed the optimizer function

```r
m3<-update(m2, start=ss, control=glmerControl(optimizer="bobyqa",optCtrl=list(maxfun=2e5)))
# model still does not converge, 11 negative eigenvalues
summary(m3)
```

```
## Generalized linear mixed model fit by maximum likelihood (Laplace
##   Approximation) [glmerMod]
##  Family: poisson  ( log )
## Formula: larvae ~ (pltstd1 + pltsqstd) * treatment + alrstd * treatment +
##     (1 | individual) + (1 | line)
##    Data: dtbdoff
## Control:
## glmerControl(optimizer = "bobyqa", optCtrl = list(maxfun = 2e+05))
##
##      AIC      BIC   logLik deviance df.resid
##  10117.3  10167.8  -5048.7  10097.3     1133
##
## Scaled residuals:
##     Min      1Q  Median      3Q     Max
## -1.5090 -0.1605 -0.0006  0.0572 17.4756
##
## Random effects:
##  Groups     Name        Variance Std.Dev.
##  line       (Intercept) 1.30203  1.141
##  individual (Intercept) 0.06812  0.261
## Number of obs: 1143, groups:  line, 1143; individual, 50
##
## Fixed effects:
##                       Estimate Std. Error z value Pr(>|z|)
## (Intercept)            2.21980    0.08811  25.193  < 2e-16 ***
## pltstd1                3.22015    0.28732  11.208  < 2e-16 ***
## pltsqstd              -6.78701    0.40736 -16.661  < 2e-16 ***
## treatmentTF            0.17477    0.10249   1.705  0.08814 .
## alrstd                 0.28953    0.07485   3.868  0.00011 ***
## pltstd1:treatmentTF   -1.15880    0.37898  -3.058  0.00223 **
## pltsqstd:treatmentTF   2.00662    0.52177   3.846  0.00012 ***
## treatmentTF:alrstd     0.04213    0.09560   0.441  0.65942
## ---
## Signif. codes:  0 '***' 0.001 '**' 0.01 '*' 0.05 '.' 0.1 ' ' 1
##
## Correlation of Fixed Effects:
##             (Intr) pltst1 pltsqs trtmTF alrstd pl1:TF plt:TF
## pltstd1     -0.489
## pltsqstd     0.576 -0.961
## treatmentTF -0.706  0.420 -0.494
```

```
## alrstd       0.089  0.032 -0.108 -0.084
## pltstd1:tTF  0.372 -0.758  0.728 -0.490 -0.025
## pltsqstd:TF -0.450  0.750 -0.780  0.591  0.083 -0.959
## trtmntTF:lr -0.082 -0.024  0.082  0.012 -0.791  0.033 -0.108
```

```
sum <- groupwiseMedian(larvae ~ plate*treatment, data = dtbdoff, conf = 0.95, R = 5000, percentile=TRUE

sum
```

```
##    plate treatment  n Median Conf.level Percentile.lower Percentile.upper
## 1      1        AL 48    6.0       0.95              4.5                8
## 2      1        TF 50    6.0       0.95              4.0                8
## 3      2        AL 47  140.0       0.95            131.0              148
## 4      2        TF 50  132.0       0.95            119.0              144
## 5      3        AL 46  196.0       0.95            191.0              208
## 6      3        TF 50  184.0       0.95            173.0              194
## 7      4        AL 46  192.0       0.95            180.0              198
## 8      4        TF 49  170.0       0.95            143.0              181
## 9      5        AL 46  117.0       0.95             97.0              134
## 10     5        TF 49   93.0       0.95             65.0              112
## 11     6        AL 45   61.0       0.95             54.0               75
## 12     6        TF 49   58.0       0.95             36.0               68
## 13     7        AL 44   37.0       0.95             28.0               43
## 14     7        TF 48   29.5       0.95             26.0               35
## 15     8        AL 41   23.0       0.95             18.0               27
## 16     8        TF 42   21.5       0.95             14.5               27
## 17     9        AL 34   13.0       0.95              6.5               19
## 18     9        TF 40   12.5       0.95             10.5               17
## 19    10        AL 27    7.0       0.95              3.0               11
## 20    10        TF 31    6.0       0.95              5.0                9
## 21    11        AL 25    5.0       0.95              3.0                6
## 22    11        TF 30    4.5       0.95              1.5                7
## 23    12        AL 23    0.0       0.95              0.0                4
## 24    12        TF 24    2.5       0.95              0.0                5
## 25    13        AL 16    0.0       0.95              0.0                2
## 26    13        TF 22    1.5       0.95              0.5                3
## 27    14        AL 14    0.0       0.95              0.0                1
## 28    14        TF 16    0.0       0.95              0.0                1
## 29    15        AL  9    0.0       0.95              0.0                2
## 30    15        TF 13    0.0       0.95              0.0                1
## 31    16        AL  8    0.0       0.95              0.0                0
## 32    16        TF 11    0.0       0.95              0.0                0
## 33    17        AL  7    0.0       0.95              0.0                0
## 34    17        TF  9    0.0       0.95              0.0                0
## 35    18        AL  6    0.0       0.95              0.0                0
## 36    18        TF  6    0.0       0.95              0.0                0
## 37    19        AL  4    0.0       0.95              0.0                0
## 38    19        TF  4    0.0       0.95              0.0                1
## 39    20        AL  4    0.0       0.95              0.0                0
## 40    20        TF  2    0.0       0.95              0.0                0
## 41    21        AL  3    0.0       0.95              0.0                0
## 42    21        TF  1    0.0       0.95              0.0                0
## 43    22        AL  2    0.0       0.95              0.0                0
## 44    22        TF  1    0.0       0.95              0.0                0
## 45    23        AL  1    0.0       0.95              0.0                0
```

```r
data2 <- subset(dtbdoff, plate == 2)
data3 <- subset(dtbdoff, plate == 3)
data4 <- subset(dtbdoff, plate == 4)

day2 <- glmer(larvae ~ pltstd1*treatment + (1|individual) + (1|line), family = poisson, data = data2)
```

```
## fixed-effect model matrix is rank deficient so dropping 2 columns / coefficients
```

```r
summary(day2)
```

```
## Generalized linear mixed model fit by maximum likelihood (Laplace
##   Approximation) [glmerMod]
##  Family: poisson  ( log )
## Formula: larvae ~ pltstd1 * treatment + (1 | individual) + (1 | line)
##    Data: data2
##
##      AIC      BIC   logLik deviance df.resid
##   1046.5   1056.8   -519.2   1038.5       93
##
## Scaled residuals:
##     Min      1Q  Median      3Q     Max
## -4.2303 -0.0594  0.0366  0.1415  0.3211
##
## Random effects:
##  Groups     Name        Variance Std.Dev.
##  line       (Intercept) 0.07049  0.2655
##  individual (Intercept) 0.04192  0.2047
## Number of obs: 97, groups:  line, 97; individual, 50
##
## Fixed effects:
##             Estimate Std. Error z value Pr(>|z|)
## (Intercept)  4.90705    0.05035  97.465   <2e-16 ***
## treatmentTF -0.13860    0.05749  -2.411   0.0159 *
## ---
## Signif. codes:  0 '***' 0.001 '**' 0.01 '*' 0.05 '.' 0.1 ' ' 1
##
## Correlation of Fixed Effects:
##             (Intr)
## treatmentTF -0.586
## fit warnings:
## fixed-effect model matrix is rank deficient so dropping 2 columns / coefficients
```

```r
day3 <- glmer(larvae ~ pltstd1*treatment + (1|individual) + (1|line), family = poisson, data = data3)
```

```
## fixed-effect model matrix is rank deficient so dropping 2 columns / coefficients
```

```
## boundary (singular) fit: see ?isSingular
```

```r
summary(day3)
```

```
## Generalized linear mixed model fit by maximum likelihood (Laplace
##   Approximation) [glmerMod]
##  Family: poisson  ( log )
## Formula: larvae ~ pltstd1 * treatment + (1 | individual) + (1 | line)
##    Data: data3
##
```

```
##      AIC      BIC   logLik deviance df.resid
##   1145.0   1155.3   -568.5   1137.0       92
##
## Scaled residuals:
##     Min      1Q  Median      3Q     Max
## -3.4736 -0.0044  0.0348  0.0705  0.1332
##
## Random effects:
##  Groups     Name        Variance Std.Dev.
##  line       (Intercept) 0.21     0.4582
##  individual (Intercept) 0.00     0.0000
## Number of obs: 96, groups:  line, 96; individual, 50
##
## Fixed effects:
##             Estimate Std. Error z value Pr(>|z|)
## (Intercept)  5.24351    0.06844  76.614   <2e-16 ***
## treatmentTF -0.21941    0.09522  -2.304   0.0212 *
## ---
## Signif. codes:  0 '***' 0.001 '**' 0.01 '*' 0.05 '.' 0.1 ' ' 1
##
## Correlation of Fixed Effects:
##             (Intr)
## treatmentTF -0.718
## fit warnings:
## fixed-effect model matrix is rank deficient so dropping 2 columns / coefficients
## convergence code: 0
## boundary (singular) fit: see ?isSingular
```

```
day4 <- glmer(larvae ~ pltstd1*treatment + (1|individual) + (1|line), family = poisson, data = data4)
```

```
## fixed-effect model matrix is rank deficient so dropping 2 columns / coefficients
## boundary (singular) fit: see ?isSingular
```

```
summary(day4)
```

```
## Generalized linear mixed model fit by maximum likelihood (Laplace
##   Approximation) [glmerMod]
##  Family: poisson  ( log )
## Formula: larvae ~ pltstd1 * treatment + (1 | individual) + (1 | line)
##    Data: data4
##
##      AIC      BIC   logLik deviance df.resid
##   1092.2   1102.4   -542.1   1084.2       91
##
## Scaled residuals:
##      Min       1Q   Median       3Q      Max
## -1.77220 -0.01448  0.04882  0.08671  0.17935
##
## Random effects:
##  Groups     Name        Variance Std.Dev.
##  line       (Intercept) 0.1965   0.4433
##  individual (Intercept) 0.0000   0.0000
## Number of obs: 95, groups:  line, 95; individual, 49
##
## Fixed effects:
```

```
##             Estimate Std. Error z value Pr(>|z|)
## (Intercept)  5.14533    0.06640  77.489   <2e-16 ***
## treatmentTF -0.19803    0.09267  -2.137   0.0326 *
## ---
## Signif. codes:  0 '***' 0.001 '**' 0.01 '*' 0.05 '.' 0.1 ' ' 1
##
## Correlation of Fixed Effects:
##             (Intr)
## treatmentTF -0.716
## fit warnings:
## fixed-effect model matrix is rank deficient so dropping 2 columns / coefficients
## convergence code: 0
## boundary (singular) fit: see ?isSingular
```

###*Check for overdispersion No overdisperison

###*Reduce data set, check it

##*Net reproductive fitness*

```
m2 <- glmer(tot_larv ~ treatment + (1|lifespan), data=dt_life_off, family='poisson')
summary(m2)
```

```
## Generalized linear mixed model fit by maximum likelihood (Laplace
##   Approximation) [glmerMod]
##  Family: poisson  ( log )
## Formula: tot_larv ~ treatment + (1 | lifespan)
##    Data: dt_life_off
##
##      AIC      BIC   logLik deviance df.resid
##   6453.3   6461.0  -3223.6   6447.3       95
##
## Scaled residuals:
##      Min       1Q   Median       3Q      Max
## -22.2957  -2.8320  -0.1355   4.7647  13.5318
##
## Random effects:
##  Groups   Name        Variance Std.Dev.
##  lifespan (Intercept) 0.3658   0.6048
## Number of obs: 98, groups:  lifespan, 21
##
## Fixed effects:
##              Estimate Std. Error z value Pr(>|z|)
## (Intercept)  6.497868   0.132244   49.14   <2e-16 ***
## treatmentTF -0.147162   0.008452  -17.41   <2e-16 ***
## ---
## Signif. codes:  0 '***' 0.001 '**' 0.01 '*' 0.05 '.' 0.1 ' ' 1
##
## Correlation of Fixed Effects:
##             (Intr)
## treatmentTF -0.028
```

```
dt_life_off[, .(mean_LRS = mean(tot_larv), median_ttllrv = median(tot_larv), sd_ttllrv = sd(tot_larv), s
```

```
##    treatment mean_LRS median_ttllrv sd_ttllrv se_ttllrv iqr_LRS
## 1:        AL 751.6667         790.5  228.3310  32.95674  215.75
## 2:        TF 690.9000         696.5  243.6565  34.45824  240.00
```

##**F1 LIFESPAN** ###Descriptive statistics and initial plots

```
##    treatment mean_lifespan median_lifespan sd_lifespan iqr_lifespan     CI
## 1:        AL      11.41667              11    4.752752            6  2.525
## 2:        AL      11.41667              11    4.752752            6 21.825
## 3:        TF      11.90000              11    4.185739            5  6.225
## 4:        TF      11.90000              11    4.185739            5 19.775
```

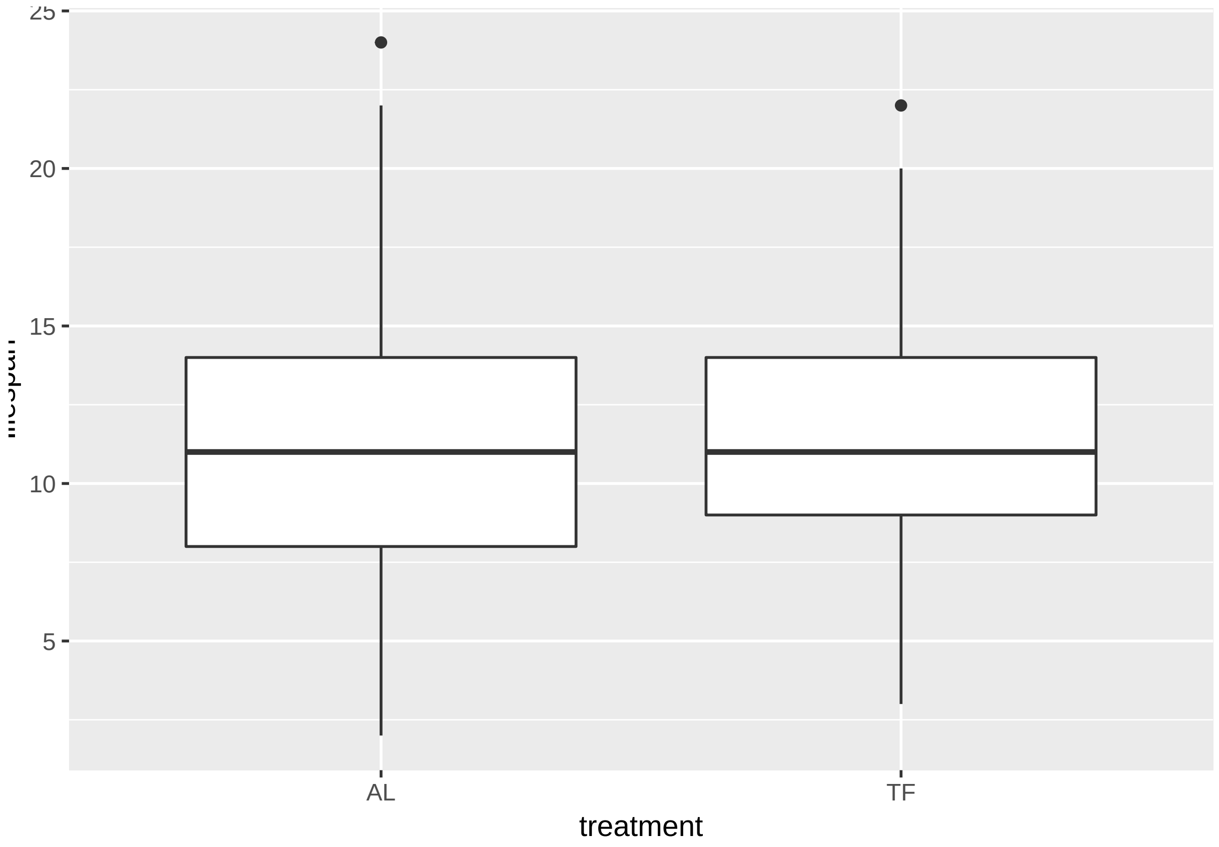

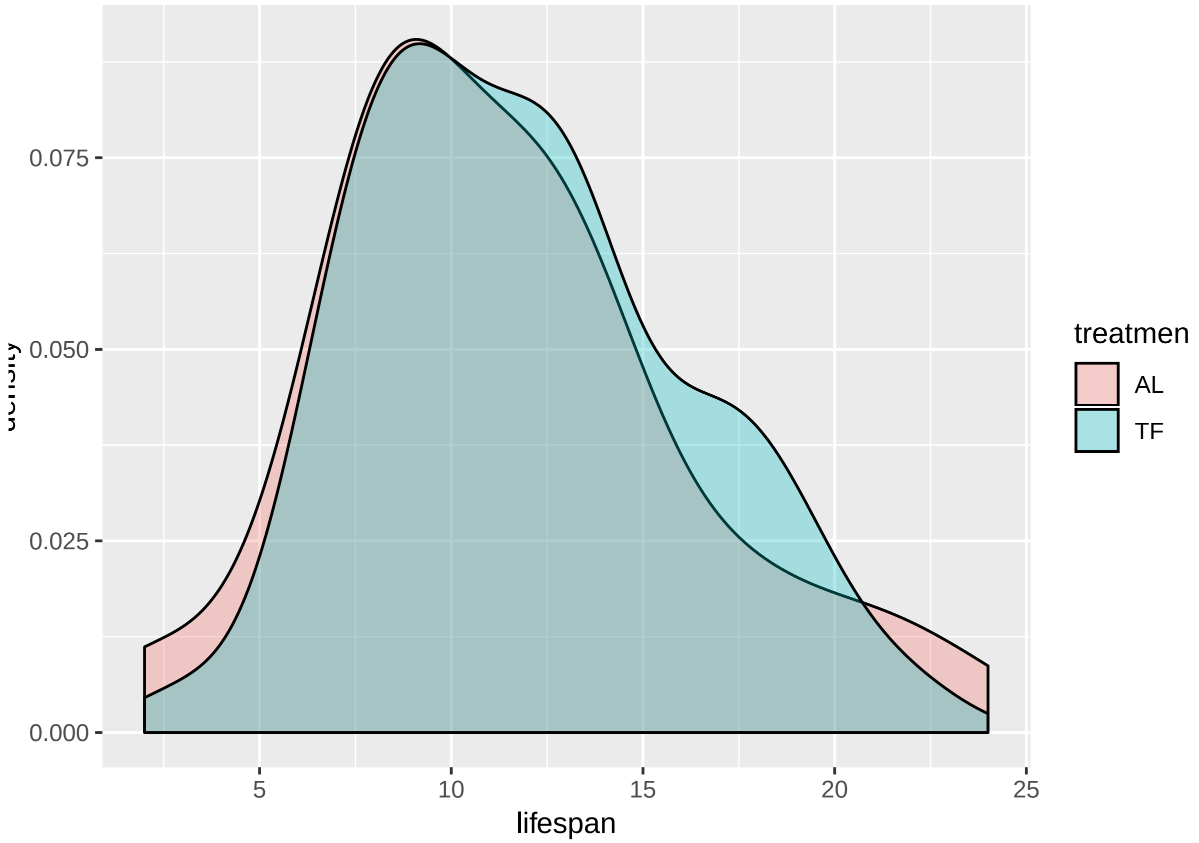


```
## Call: survfit(formula = Surv(lifespan, Event) ~ treatment, data = dt_life_off)
##
##                 treatment=AL
##  time n.risk n.event survival std.err lower 95% CI upper 95% CI
##     2     48       2   0.9583  0.0288       0.9034        1.000
##     5     46       1   0.9375  0.0349       0.8715        1.000
##     6     45       1   0.9167  0.0399       0.8417        0.998
##     7     44       3   0.8542  0.0509       0.7599        0.960
##     8     41       7   0.7083  0.0656       0.5907        0.849
##     9     34       7   0.5625  0.0716       0.4383        0.722
##    10     27       2   0.5208  0.0721       0.3971        0.683
##    11     25       2   0.4792  0.0721       0.3568        0.644
##    12     23       7   0.3333  0.0680       0.2234        0.497
##    13     16       2   0.2917  0.0656       0.1877        0.453
##    14     14       5   0.1875  0.0563       0.1041        0.338
##    15      9       1   0.1667  0.0538       0.0885        0.314
##    16      8       1   0.1458  0.0509       0.0735        0.289
##    17      7       1   0.1250  0.0477       0.0591        0.264
##    18      6       2   0.0833  0.0399       0.0326        0.213
##    20      4       1   0.0625  0.0349       0.0209        0.187
##    21      3       1   0.0417  0.0288       0.0107        0.162
##    22      2       1   0.0208  0.0206       0.0030        0.145
##    24      1       1   0.0000     NaN           NA           NA
##
##                 treatment=TF
##  time n.risk n.event survival std.err lower 95% CI upper 95% CI
##     3     50       1     0.98  0.0198      0.94195        1.000
```

```
##     6     49       1     0.96  0.0277      0.90719        1.000
##     7     48       6     0.84  0.0518      0.74429        0.948
##     8     42       2     0.80  0.0566      0.69647        0.919
##     9     40       9     0.62  0.0686      0.49906        0.770
##    10     31       1     0.60  0.0693      0.47848        0.752
##    11     30       6     0.48  0.0707      0.35971        0.641
##    12     24       2     0.44  0.0702      0.32185        0.602
##    13     22       7     0.30  0.0648      0.19644        0.458
##    14     15       3     0.24  0.0604      0.14655        0.393
##    15     12       1     0.22  0.0586      0.13054        0.371
##    16     11       2     0.18  0.0543      0.09962        0.325
##    17      9       3     0.12  0.0460      0.05665        0.254
##    18      6       2     0.08  0.0384      0.03125        0.205
##    19      4       2     0.04  0.0277      0.01029        0.156
##    20      2       1     0.02  0.0198      0.00287        0.139
##    22      1       1     0.00     NaN           NA           NA
```

```
## Call: survfit(formula = Surv(lifespan, Event) ~ treatment, data = dt_life_off)
##
##                 n events median 0.95LCL 0.95UCL
## treatment=AL 48     48     11       9      12
## treatment=TF 50     50     11       9      13
```

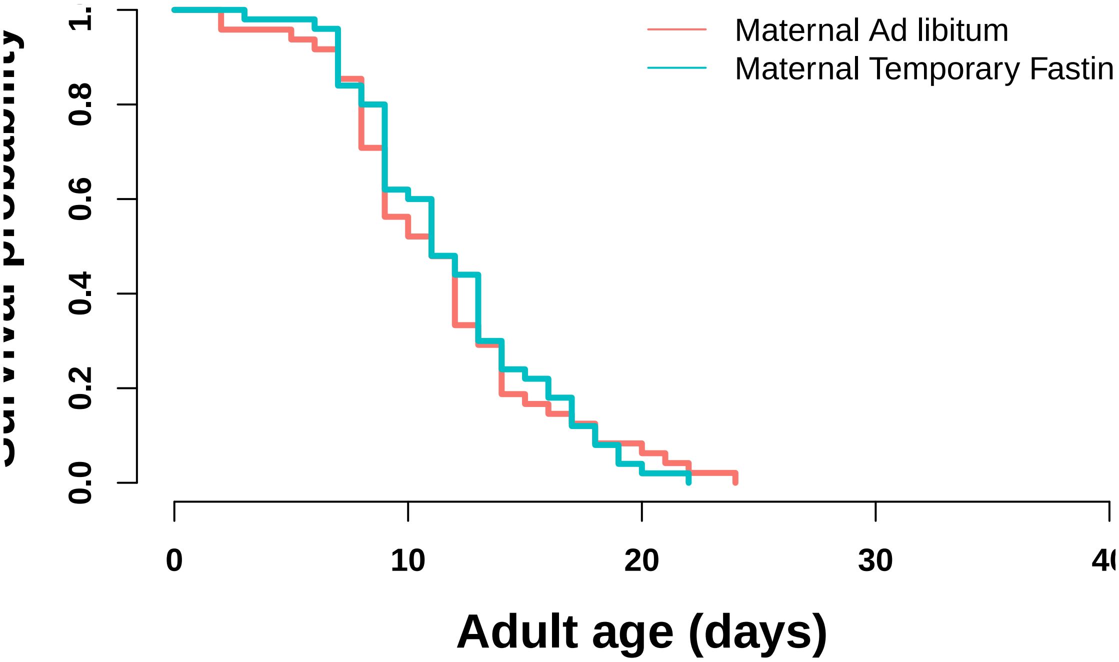


###Statistical model There is no difference in lifespan between offspring of mothers in either treatment (both median lifespan 11 days 9-13 days (95% CI)).

```
#ANALYSIS USING COXPH IN SURVIVAL package
m_off_life <- coxph(Surv(lifespan,Event)~treatment ,data=dt_life_off)
summary(m_off_life)
```

```
## Call:
## coxph(formula = Surv(lifespan, Event) ~ treatment, data = dt_life_off)
##
##   n= 98, number of events= 98
```

```
##
##                coef exp(coef) se(coef)      z Pr(>|z|)
## treatmentTF -0.0196    0.9806   0.2045 -0.096    0.924
##
##             exp(coef) exp(-coef) lower .95 upper .95
## treatmentTF    0.9806       1.02    0.6567     1.464
##
## Concordance= 0.521  (se = 0.032 )
## Likelihood ratio test= 0.01  on 1 df,   p=0.9
## Wald test            = 0.01  on 1 df,   p=0.9
## Score (logrank) test = 0.01  on 1 df,   p=0.9
```

```
#check assumption of the ph model
temp <- cox.zph(m_off_life)
temp
```

```
##               rho chisq     p
## treatmentTF 0.112  1.24 0.266
```

```
plot(temp)
```

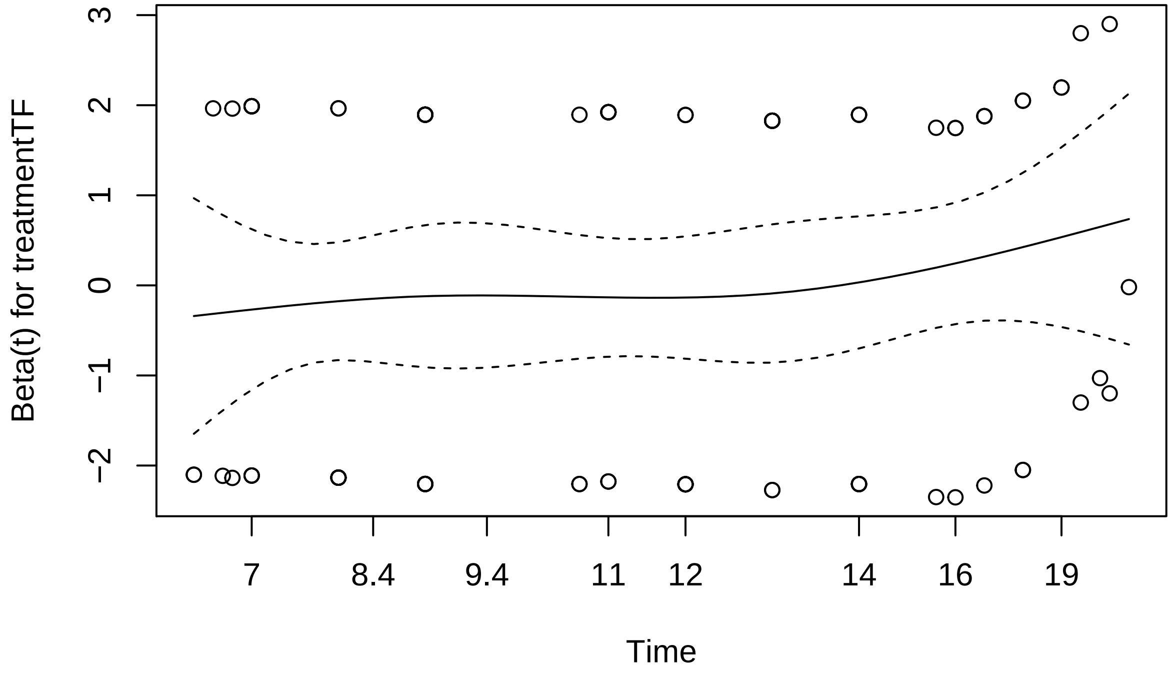


##**F1 HEAT SHOCK** ####*data uploaded and new variable created.

###Statistical model There is no difference between offspring with mothers from the Temporary Fasting treatment and those with mothers from ad libitum treatment (Wald Z=0.892, p >0.372) and no interaction with time (see model table below). There was a significant lower proportion of unaffected individuals with time (see table below).

```
m_f1hs <- glmer(response ~ treatment*time  + (1|plt_combo), data= dt_F1hs, family="binomial")
```

```
## Warning in checkConv(attr(opt, "derivs"), opt$par, ctrl =
## control$checkConv, : unable to evaluate scaled gradient

## Warning in checkConv(attr(opt, "derivs"), opt$par, ctrl =
## control$checkConv, : Model failed to converge: degenerate Hessian with 1
```

```
## negative eigenvalues
```

```r
summary(m_f1hs)
```

```
## Warning in vcov.merMod(object, use.hessian = use.hessian): variance-covariance matrix computed from f
## not positive definite or contains NA values: falling back to var-cov estimated from RX

## Warning in vcov.merMod(object, correlation = correlation, sigm = sig): variance-covariance matrix com
## not positive definite or contains NA values: falling back to var-cov estimated from RX

## Generalized linear mixed model fit by maximum likelihood (Laplace
##   Approximation) [glmerMod]
##  Family: binomial  ( logit )
## Formula: response ~ treatment * time + (1 | plt_combo)
##    Data: dt_F1hs
##
##      AIC      BIC   logLik deviance df.resid
##    266.5    303.0   -118.3    236.5       69
##
## Scaled residuals:
##      Min       1Q   Median       3Q      Max
## -2.16064 -0.64775 -0.09825  0.69978  2.39118
##
## Random effects:
##  Groups    Name        Variance Std.Dev.
##  plt_combo (Intercept) 0.07933  0.2816
## Number of obs: 84, groups:  plt_combo, 6
##
## Fixed effects:
##                       Estimate Std. Error z value Pr(>|z|)
## (Intercept)            2.41681    0.48521   4.981 6.33e-07 ***
## treatmenttf           -0.54179    0.60725  -0.892 0.372282
## time3                 -1.95591    0.54239  -3.606 0.000311 ***
## time4                 -3.33401    0.55318  -6.027 1.67e-09 ***
## time5                 -4.13917    0.59655  -6.939 3.96e-12 ***
## time6                 -5.06650    0.70335  -7.203 5.87e-13 ***
## time7                 -6.50973    1.12056  -5.809 6.27e-09 ***
## time24                -0.82413    0.58699  -1.404 0.160327
## treatmenttf:time3     -0.09837    0.71404  -0.138 0.890422
## treatmenttf:time4      0.35792    0.73672   0.486 0.627086
## treatmenttf:time5      0.87007    0.77965   1.116 0.264429
## treatmenttf:time6     -0.19991    1.07996  -0.185 0.853143
## treatmenttf:time7    -14.00681 1433.92906  -0.010 0.992206
## treatmenttf:time24     0.66901    0.79040   0.846 0.397315
## ---
## Signif. codes:  0 '***' 0.001 '**' 0.01 '*' 0.05 '.' 0.1 ' ' 1
## convergence code: 0
## unable to evaluate scaled gradient
## Model failed to converge: degenerate  Hessian with 1 negative eigenvalues
```

##**F1 DEVELOPMENT TIME** ####*Data upload, time conversion, and data check

###Descriptive statistics and initial plots

```r
#Table of descriptive stats

dtdev[ , .(Mean = mean(dev_time), Median = median(dev_time), St_Dev = sd(dev_time), se_dev = sd(dev_time
```

```
##    treatment     Mean Median   St_Dev    se_dev IQR
## 1:        al 67.22348   67.0 2.483007 0.2161180 3.0
## 2:        tf 68.58333   68.5 2.499746 0.2175749 2.5
```

```r
### basic plot
ggplot(dtdev, aes(x= treatment, y = dev_time, fill=treatment)) +
  geom_boxplot()+
  theme_bw()
```

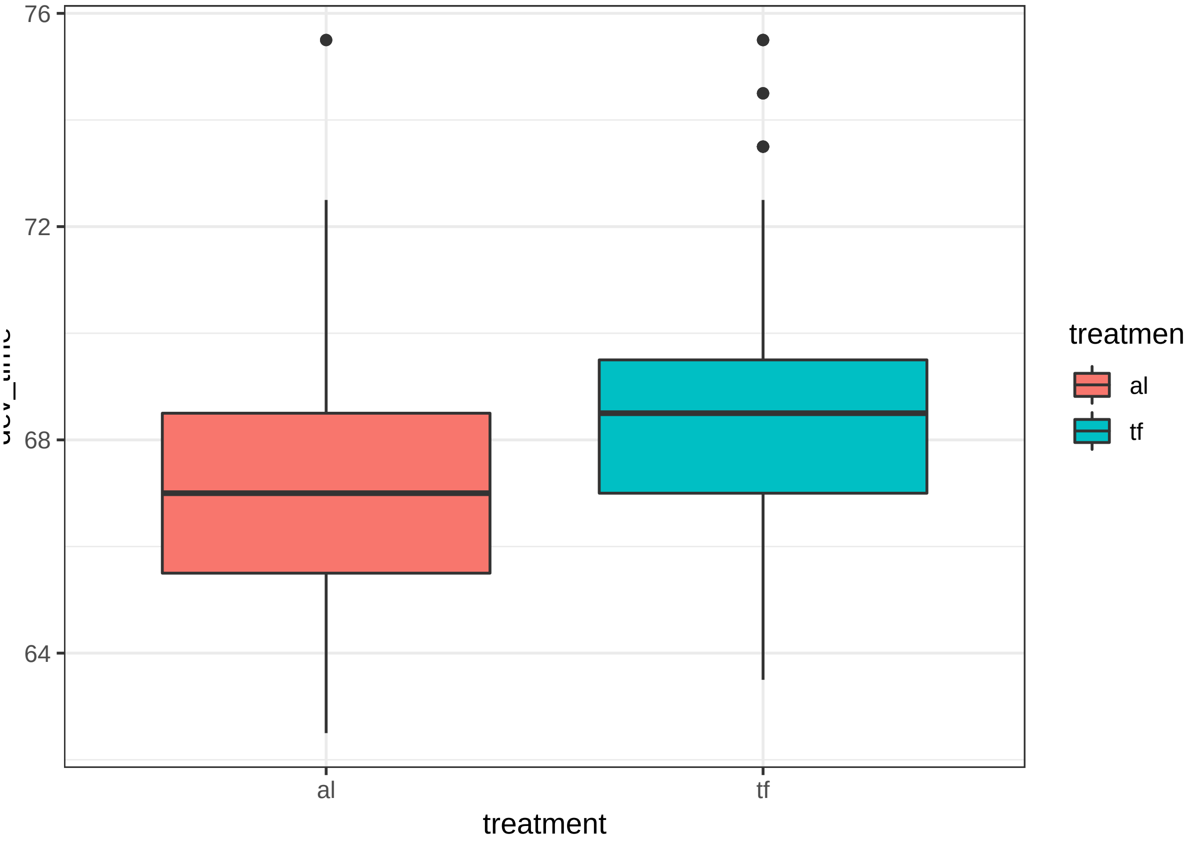


###Development analysis

Initial model with maternal plate as a 'block' factor accounted for zero variance and was dropped from the glmm model, reducing the model to a basic glm. The glm model runs fine, but residuals are not normal. This is likely due to a few outliers. Outliers were removed from the analysis and model rerun. Residuals look fine from this model. Results are qualitatively similar with treatment having a siginficant effect on development time without the outliers.

Offspring from mothers in the Temporary Fasting treatment took signficantly longer to develop (68.6 +/- 2.50 hours) than offspring from mothers in the ad libitum treatment (67.2 +/- 2.48 hrs; t= -3.07, p = 0.002).

```r
#####Analysis
m0 <- lmer(dev_time ~ treatment + (1|plate), data = dtdev)
```

```
## boundary (singular) fit: see ?isSingular
```

```r
summary(m0)
```

```
## Linear mixed model fit by REML ['lmerMod']
## Formula: dev_time ~ treatment + (1 | plate)
##    Data: dtdev
```

```
##
## REML criterion at convergence: 1231.6
##
## Scaled residuals:
##     Min      1Q  Median      3Q     Max
## -2.0404 -0.6918 -0.0897  0.5124  3.3220
##
## Random effects:
##  Groups   Name        Variance Std.Dev.
##  plate    (Intercept) 0.000    0.000
##  Residual             6.207    2.491
## Number of obs: 264, groups:  plate, 3
##
## Fixed effects:
##             Estimate Std. Error t value
## (Intercept)  67.2235     0.2168 310.003
## treatmenttf   1.3598     0.3067   4.434
##
## Correlation of Fixed Effects:
##             (Intr)
## treatmenttf -0.707
## convergence code: 0
## boundary (singular) fit: see ?isSingular
```

```
plot(m0)
```

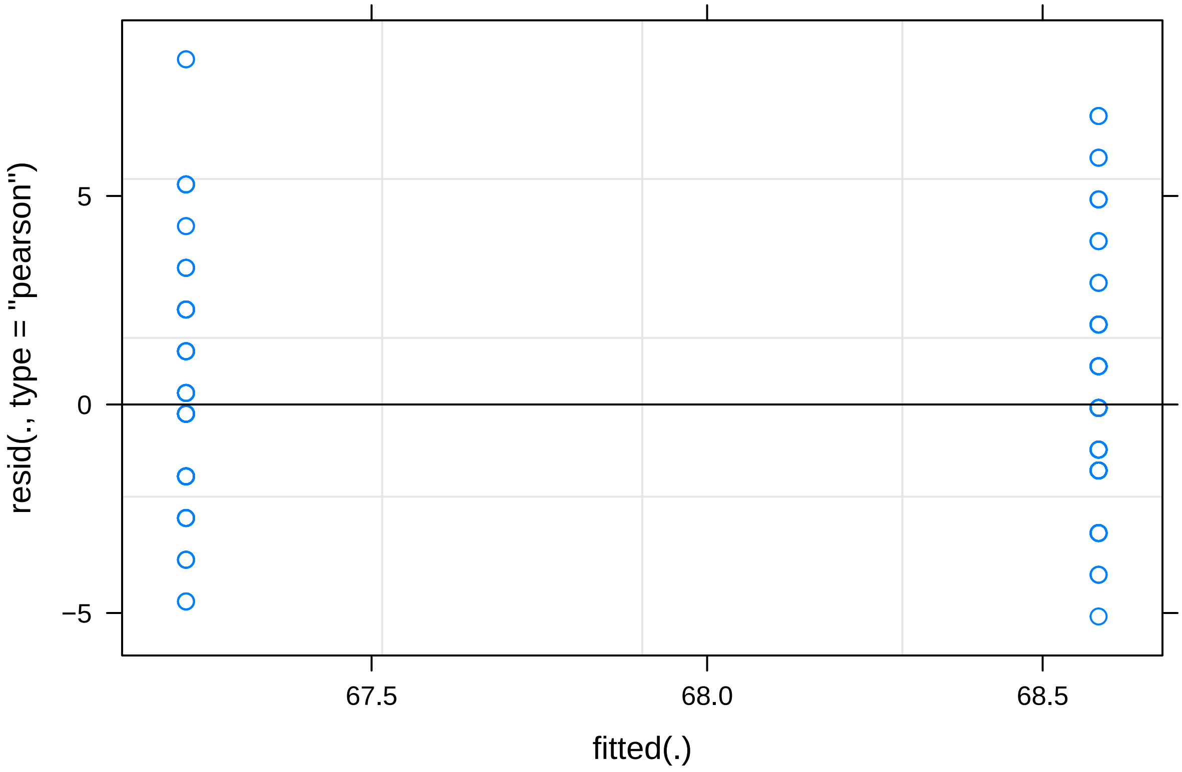


```
#plate accounts for zero variance, removed from model

m1 <- glm(dev_time ~ treatment, data = dtdev)
summary(m1)
```

```
##
## Call:
## glm(formula = dev_time ~ treatment, data = dtdev)
##
## Deviance Residuals:
##     Min       1Q   Median       3Q      Max
## -5.0833  -1.7235  -0.2235   1.2765   8.2765
##
## Coefficients:
##             Estimate Std. Error t value Pr(>|t|)
## (Intercept)  67.2235     0.2168 310.003  < 2e-16 ***
## treatmenttf   1.3598     0.3067   4.434 1.36e-05 ***
## ---
## Signif. codes:  0 '***' 0.001 '**' 0.01 '*' 0.05 '.' 0.1 ' ' 1
##
## (Dispersion parameter for gaussian family taken to be 6.207025)
##
##     Null deviance: 1748.3  on 263  degrees of freedom
## Residual deviance: 1626.2  on 262  degrees of freedom
## AIC: 1235.2
##
## Number of Fisher Scoring iterations: 2
```

```r
plot(m1)
```

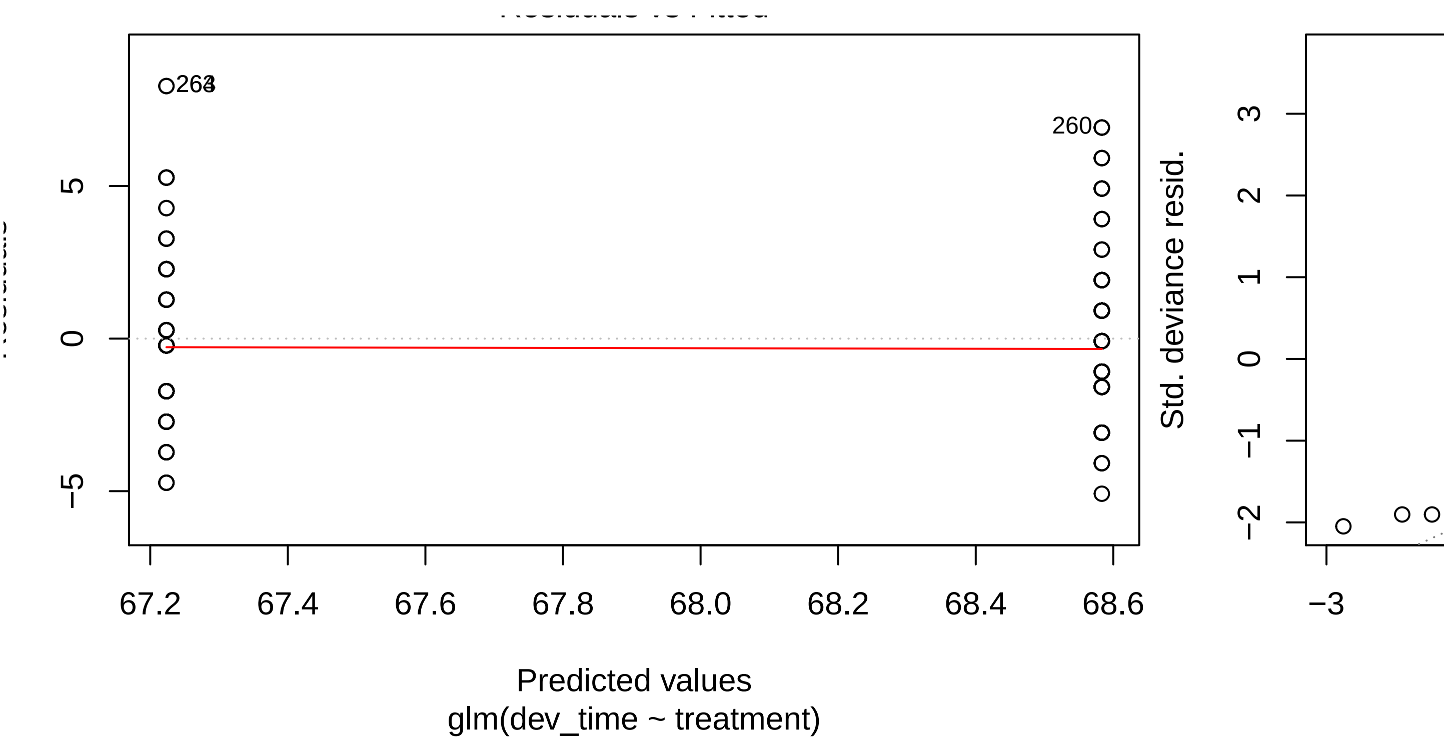


```
## hat values (leverages) are all = 0.007575758
##  and there are no factor predictors; no plot no. 5
```

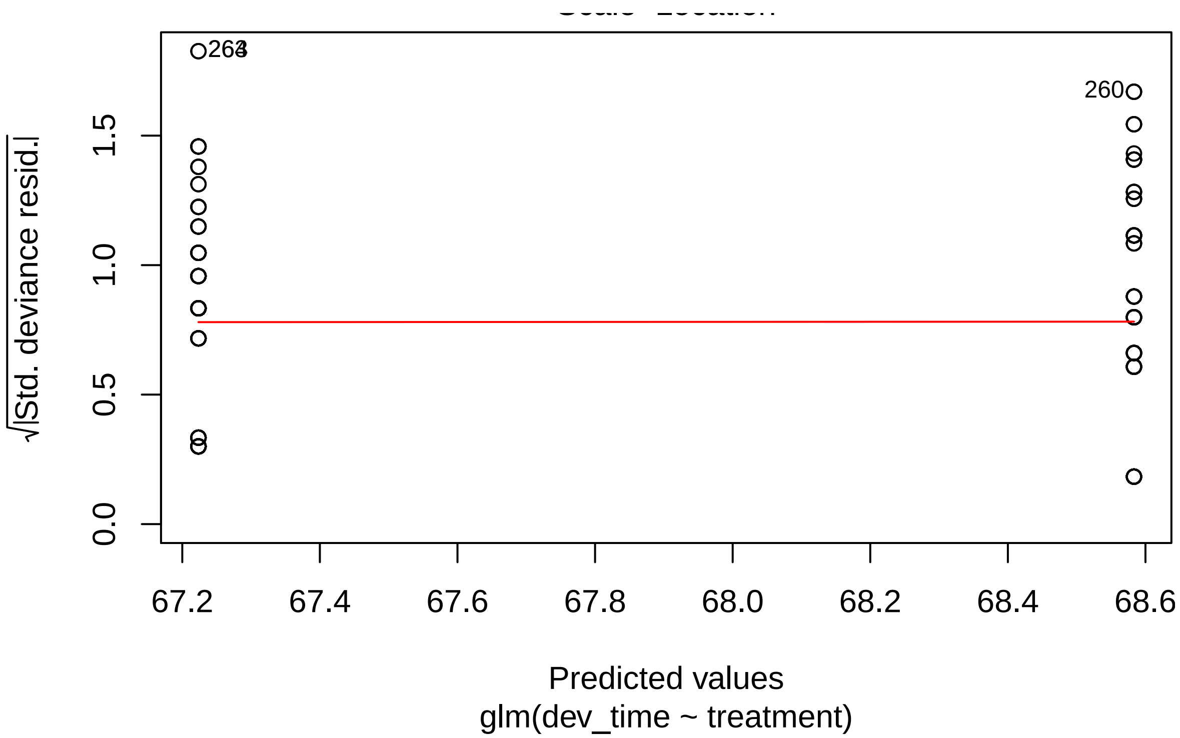


```r
#Analysis without outliers

dev1 <- dtdev[-c(93,184,201),]
m4 <- glm(size ~ treatment, data = dev1)
summary(m4)
```

```
##
## Call:
## glm(formula = size ~ treatment, data = dev1)
##
## Deviance Residuals:
##        Min          1Q      Median          3Q         Max
## -0.0059711  -0.0017131  -0.0001651   0.0016736   0.0065906
##
## Coefficients:
##               Estimate Std. Error t value Pr(>|t|)
## (Intercept)  0.0302481  0.0002215 136.571   <2e-16 ***
## treatmenttf -0.0009547  0.0003114  -3.066   0.0024 **
## ---
## Signif. codes:  0 '***' 0.001 '**' 0.01 '*' 0.05 '.' 0.1 ' ' 1
##
## (Dispersion parameter for gaussian family taken to be 6.328038e-06)
##
##     Null deviance: 0.0016984  on 260  degrees of freedom
## Residual deviance: 0.0016390  on 259  degrees of freedom
## AIC: -2379.6
##
## Number of Fisher Scoring iterations: 2
```

```r
plot(m4)
```

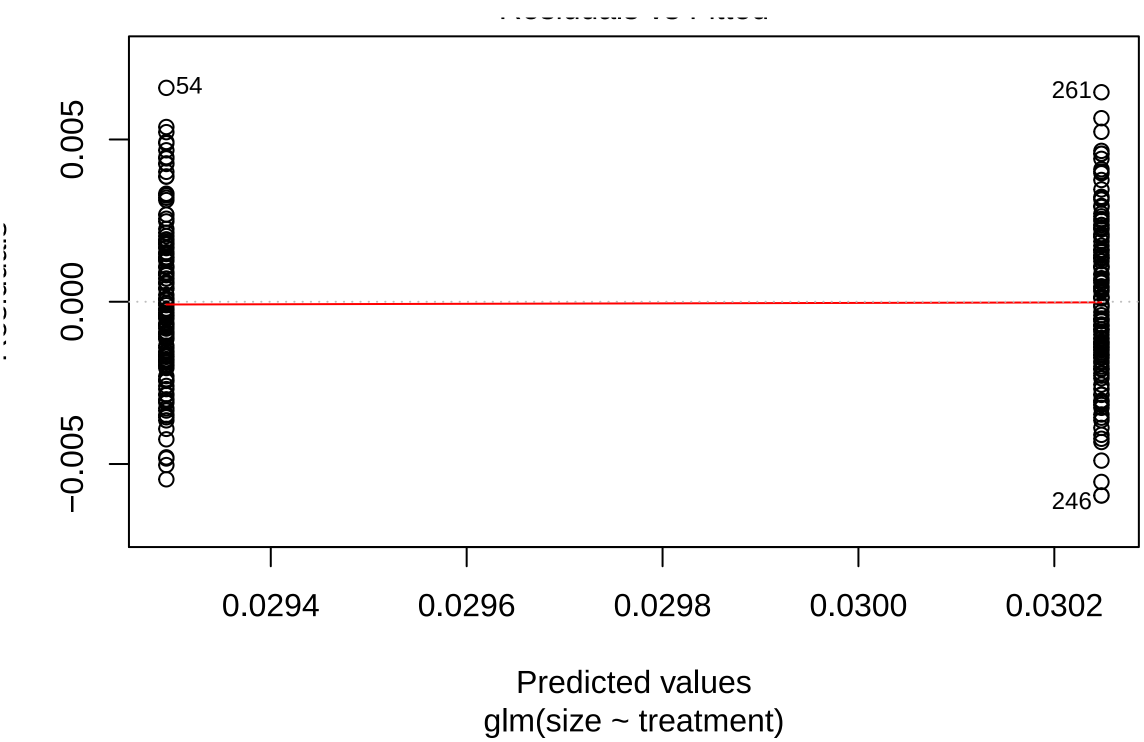

54
261
246
0.005
0.000
−0.005
0.0294
0.0296
0.0298
0.0300
0.0302
Predicted values
glm(size ~ treatment)


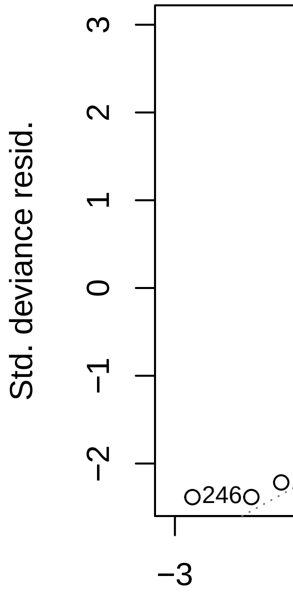

Std. deviance resid.
3
2
1
0
−1
−2
246
−3


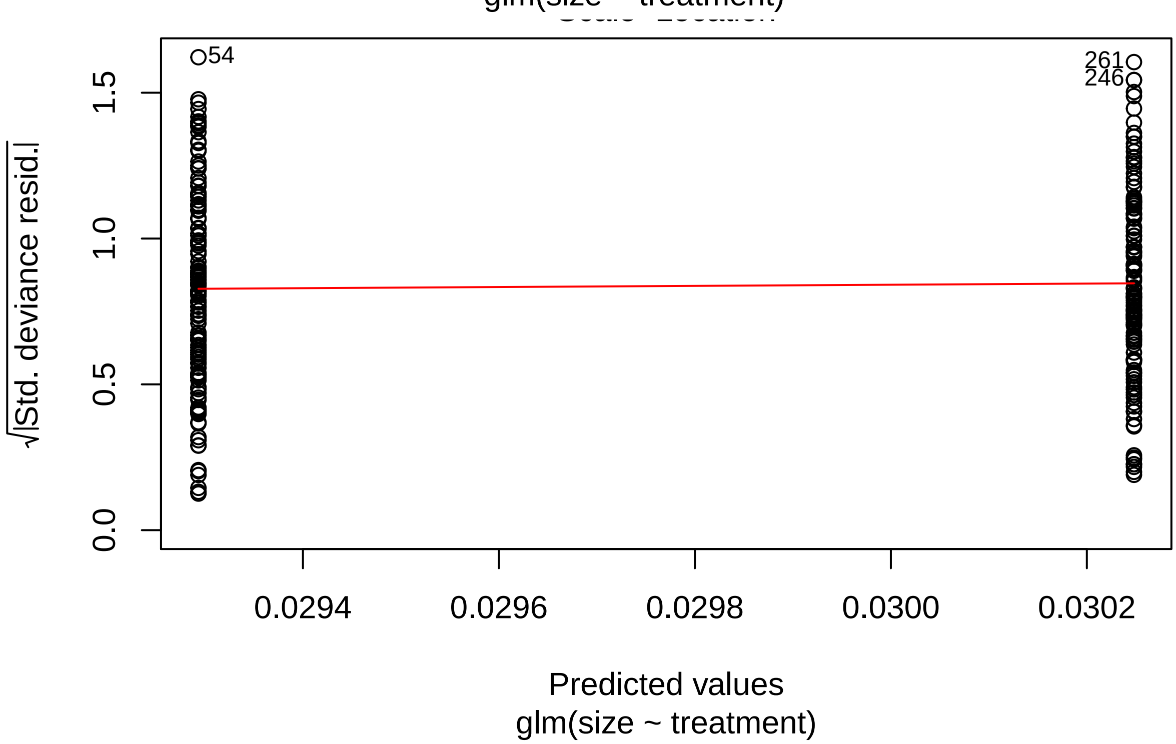

54
261
246
1.5
1.0
0.5
0.0
√|Std. deviance resid.|
0.0294
0.0296
0.0298
0.0300
0.0302
Predicted values
glm(size ~ treatment)

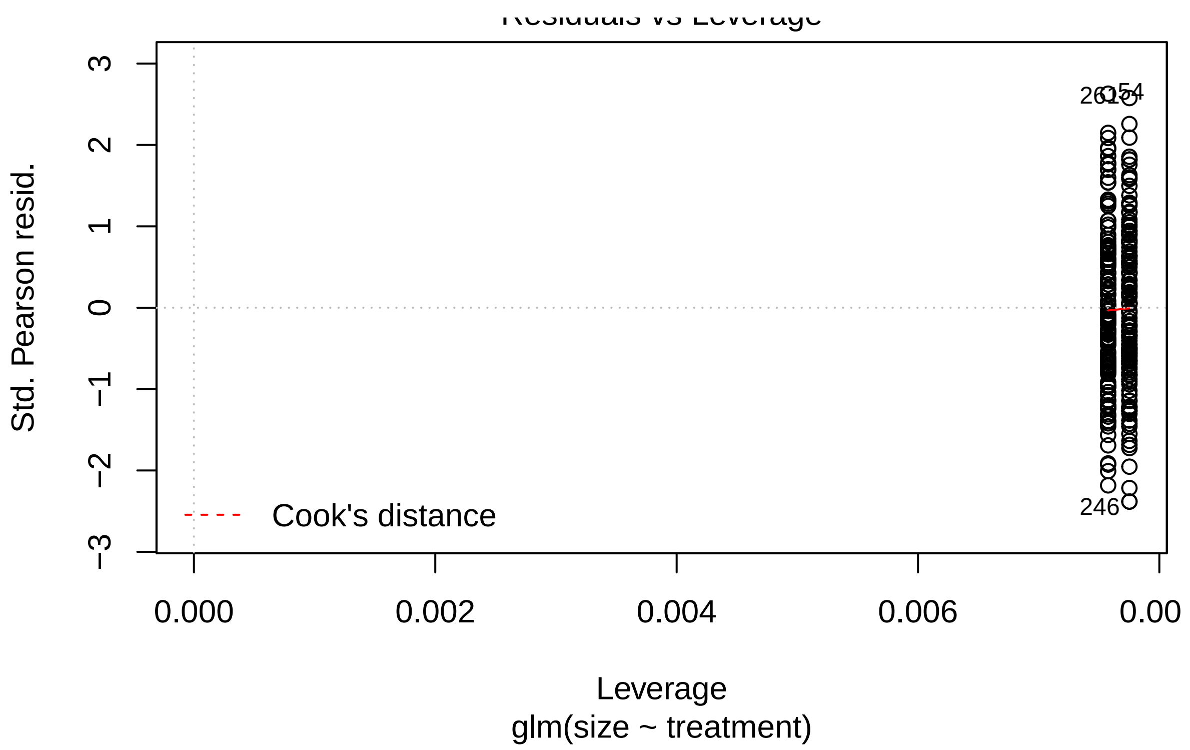


##**F1 BODY SIZE** ###Descriptive statistics and initial plot

```
#descriptive statistics
dtdev[ , .(Mean = mean(size), Median = median(size), St_Dev = sd(size), se_size = sd(size)/sqrt(length(
```

```
##    treatment       Mean    Median      St_Dev      se_size        IQR
## 1:        al 0.03020696 0.0301390 0.002836712 0.0002469041 0.00347850
## 2:        tf 0.02929337 0.0290465 0.002486697 0.0002164392 0.00342975
```

```
#plot
ggplot(dtdev, aes(x=treatment, y= size, fill= treatment))+
  geom_boxplot()+
  theme_bw()
```

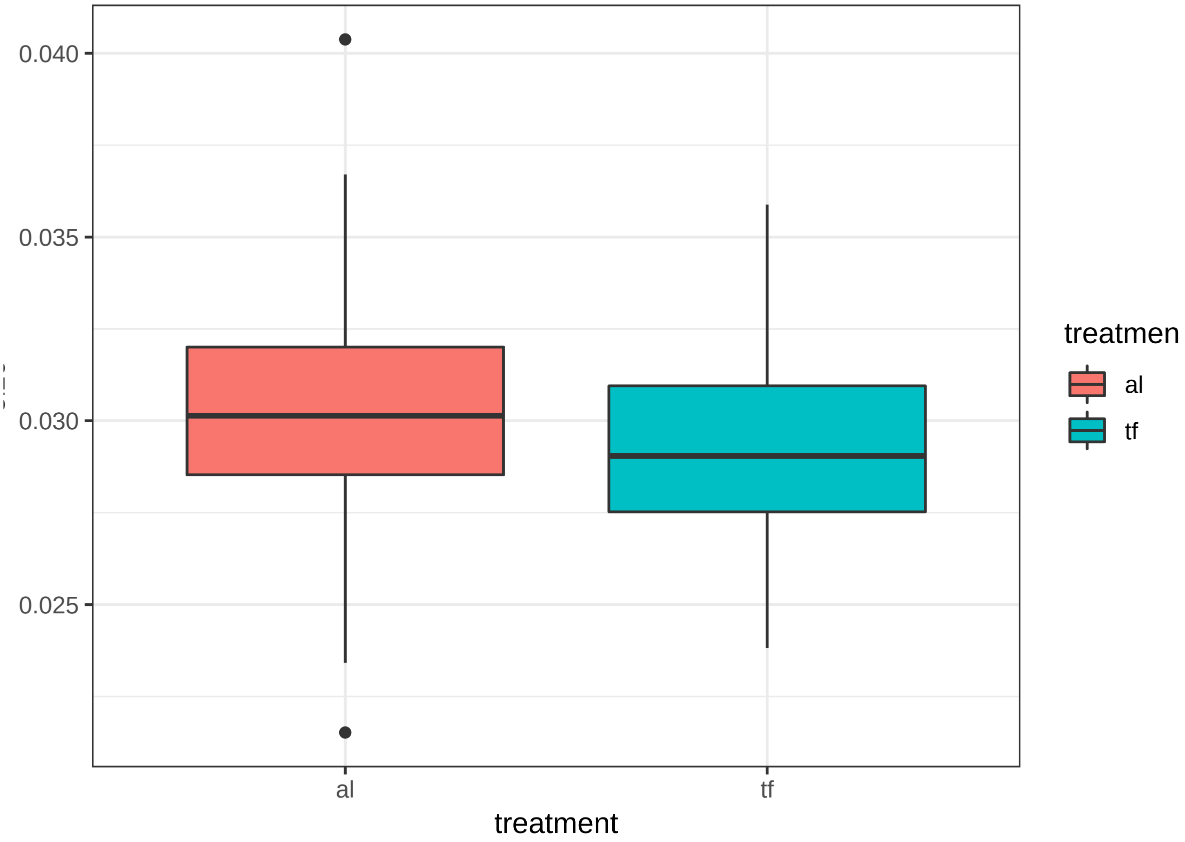


###Body size analysis The glmm with maternal (laying) plate as a random factor accounts for basically 0 variance and was dropped from the model.

Offspring from female in the Temporary Fasting experiment were signficantly smaller (0.0293 +/- 0.0025 sq mm) than those with mothers in the ad libitium treatment (0.0302 +/- 0.0028; t= 2.78, p < 0.0058).

```
m_size <- lmer(size ~ treatment + (1|plate), data = dtdev)
summary(m_size)
```

```
## Linear mixed model fit by REML ['lmerMod']
## Formula: size ~ treatment + (1 | plate)
##    Data: dtdev
##
## REML criterion at convergence: -2352.8
##
## Scaled residuals:
##     Min      1Q  Median      3Q     Max
## -3.2511 -0.6470 -0.0717  0.6465  3.7558
##
## Random effects:
##  Groups   Name        Variance  Std.Dev.
##  plate    (Intercept) 6.658e-08 0.000258
##  Residual             7.070e-06 0.002659
## Number of obs: 264, groups:  plate, 3
##
## Fixed effects:
##              Estimate Std. Error t value
## (Intercept)  0.0302018  0.0002753 109.692
```

```
## treatmenttf -0.0009180  0.0003286  -2.794
##
## Correlation of Fixed Effects:
##             (Intr)
## treatmenttf -0.594
```

```
coefs <- data.frame(coef(summary(m_size)))
coefs$p.z <- 2 * (1 - pnorm(abs(coefs$t.value)))
coefs
```

```
##                 Estimate   Std..Error  t.value        p.z
## (Intercept)  0.0302018423 0.0002753342 109.6916 0.00000000
## treatmenttf -0.0009180112 0.0003285652  -2.7940 0.00520605
```

```
plot(m_size)
```

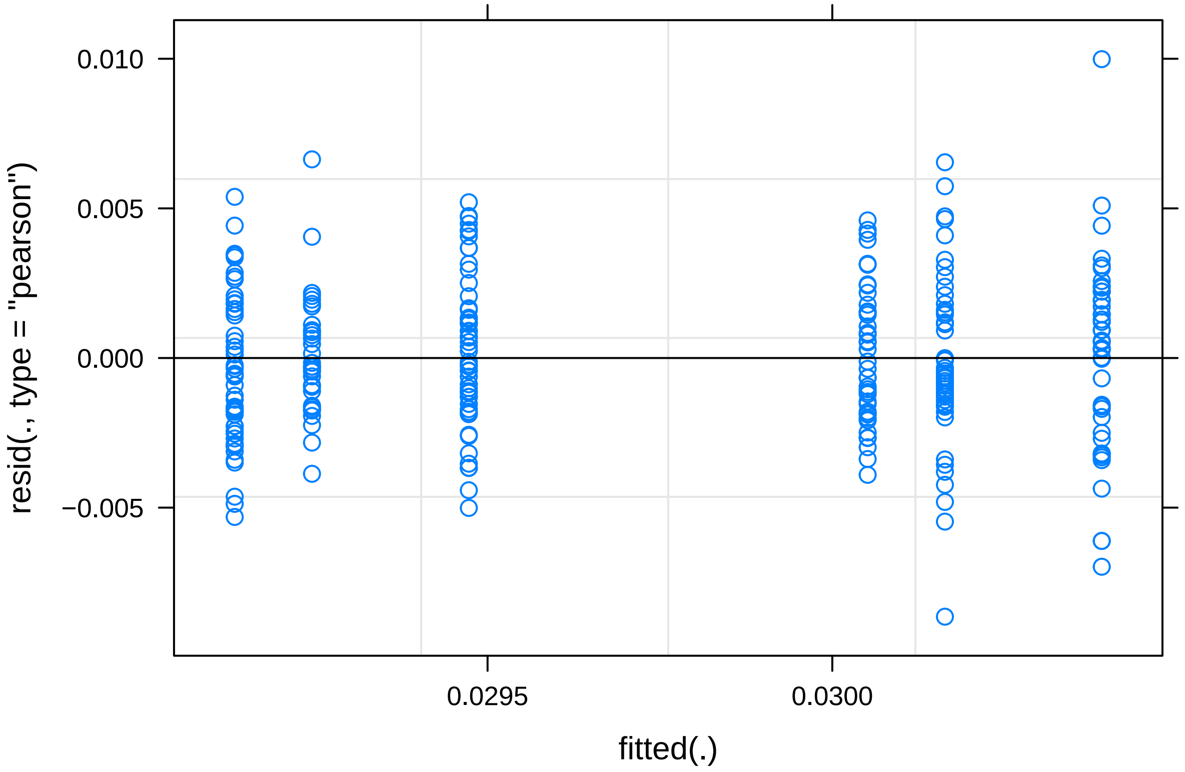


```
#plate accounts for basically zero variance, dropped from model

m2_size <- glm(size ~ treatment, data = dtdev)
summary(m2_size)
```

```
##
## Call:
## glm(formula = size ~ treatment, data = dtdev)
##
## Deviance Residuals:
##        Min          1Q      Median          3Q         Max
## -0.0086880  -0.0017141  -0.0001672   0.0016744   0.0101700
##
## Coefficients:
##                 Estimate Std. Error t value Pr(>|t|)
```

```
## (Intercept)  0.0302070  0.0002322 130.106  < 2e-16 ***
## treatmenttf -0.0009136  0.0003283  -2.782  0.00579 **
## ---
## Signif. codes:  0 '***' 0.001 '**' 0.01 '*' 0.05 '.' 0.1 ' ' 1
##
## (Dispersion parameter for gaussian family taken to be 7.115298e-06)
##
##     Null deviance: 0.0019193  on 263  degrees of freedom
## Residual deviance: 0.0018642  on 262  degrees of freedom
## AIC: -2376.1
##
## Number of Fisher Scoring iterations: 2
```

```r
plot(m2_size)
```

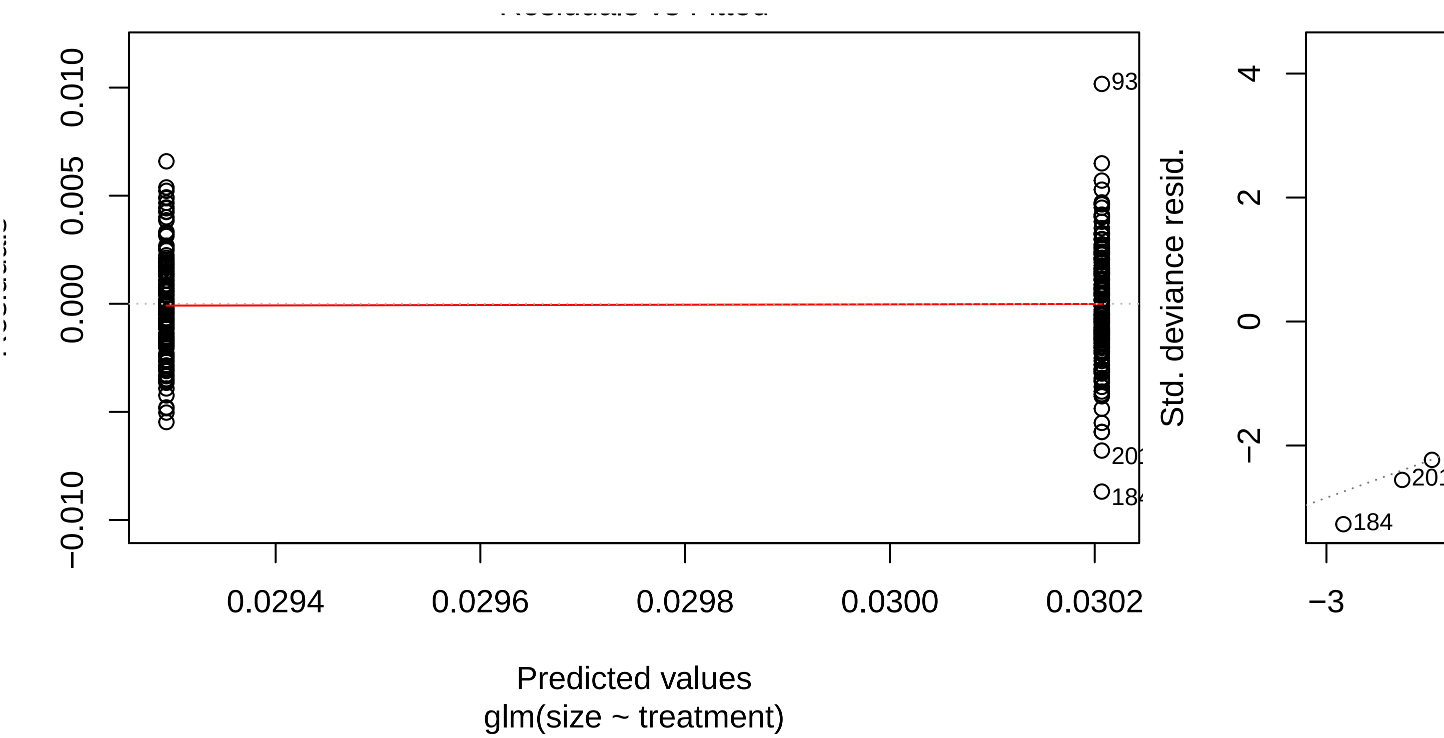


```
## hat values (leverages) are all = 0.007575758
##  and there are no factor predictors; no plot no. 5
```

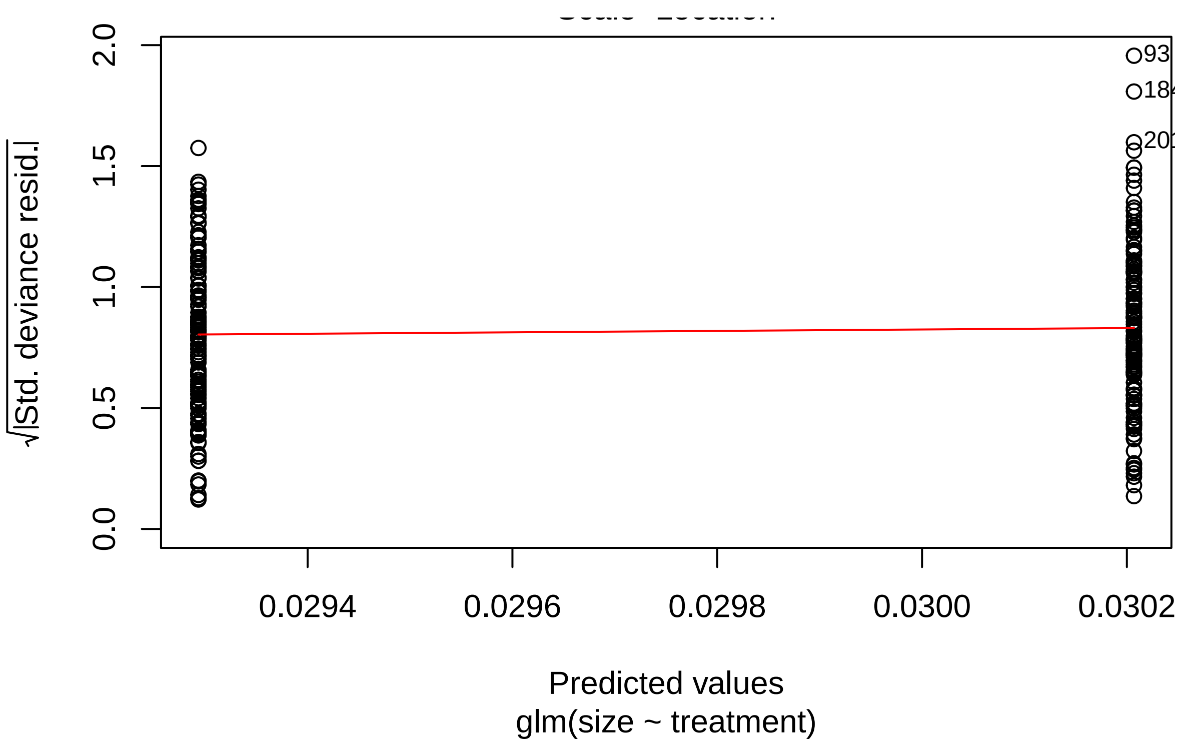

√|Std. deviance resid.|
0.0
0.5
1.0
1.5
2.0
0.0294
0.0296
0.0298
0.0300
0.0302
Predicted values
glm(size ~ treatment)
